# PowerCHORD: constructing optimal experimental designs for biological rhythm discovery

**DOI:** 10.1101/2024.05.19.594858

**Authors:** Turner Silverthorne, Matthew Carlucci, Arturas Petronis, Adam R Stinchcombe

## Abstract

Equally spaced temporal sampling is the standard protocol for the study of biological rhythms. These equispaced designs perform well when calibrated to an oscillator’s period yet can have systematic detection biases when applied to rhythms of unknown periodicity. Here, we present a broadly-applicable set of computational methods for seeking optimal measurement schedules for rhythm detection. Our PowerCHORD methods generate experimental designs by maximizing a closed-form expression for the statistical power of the cosinor model using a black-box optimization method (differential evolution), a brute-force search, or mixed-integer conic programming. Application of these three methods showed numerically that they improve upon equispaced designs under many experimental contexts. Our numerical results also revealed an intuitive approach for achieving optimal power for simultaneous investigation of circadian, circalunar, and circannual rhythms. Our findings suggest that timing optimization is an effective yet under-explored tool for improving biological rhythm discovery.

## 1 Introduction

Biological rhythms serve essential roles in living systems and arise from diverse mechanisms on scales ranging from individual cells to entire populations [1–4]. While familiar examples such as circadian rhythms, somitogenesis, and the cell-cycle have been studied for decades [5–7], new rhythms and layers of temporal organization continue to be discovered [8–12]. In the context of circadian studies, rhythm discovery has been facilitated in part by improvements to statistical methods [13–16], leading to evidence of new rhythmicity in existing datasets [17]. These improvements are now a major focus of rhythm discovery guidelines [18, 19], however the dependence of statistical performance on the underlying experimental design is less thoroughly explored. Hence, many studies make use of equispaced temporal sampling as the *de facto* design choice.

The preferred status of equispaced designs is justifiable by the strong assumptions of circadian studies. In particular, if the period of an oscillator is known ahead of time, designs with measurements equispaced along the oscillator’s cycle achieve statistically optimal performance [20, 21]. Yet, in more general contexts, biological oscillations are known to occur with periods ranging from milliseconds to years and a study may need to consider potential cycles across these vast timescales. Equispaced collection along only one cycle may be unreliable for rhythm detection due to statistical power variability at nominal periods and acrophases (Supp. Fig. 1). Given the inadequacy of equispaced designs for exploring multiple cycles, there is a need for methods that enable optimal detection of cycles with unknown periodicity.

**Figure 1:**
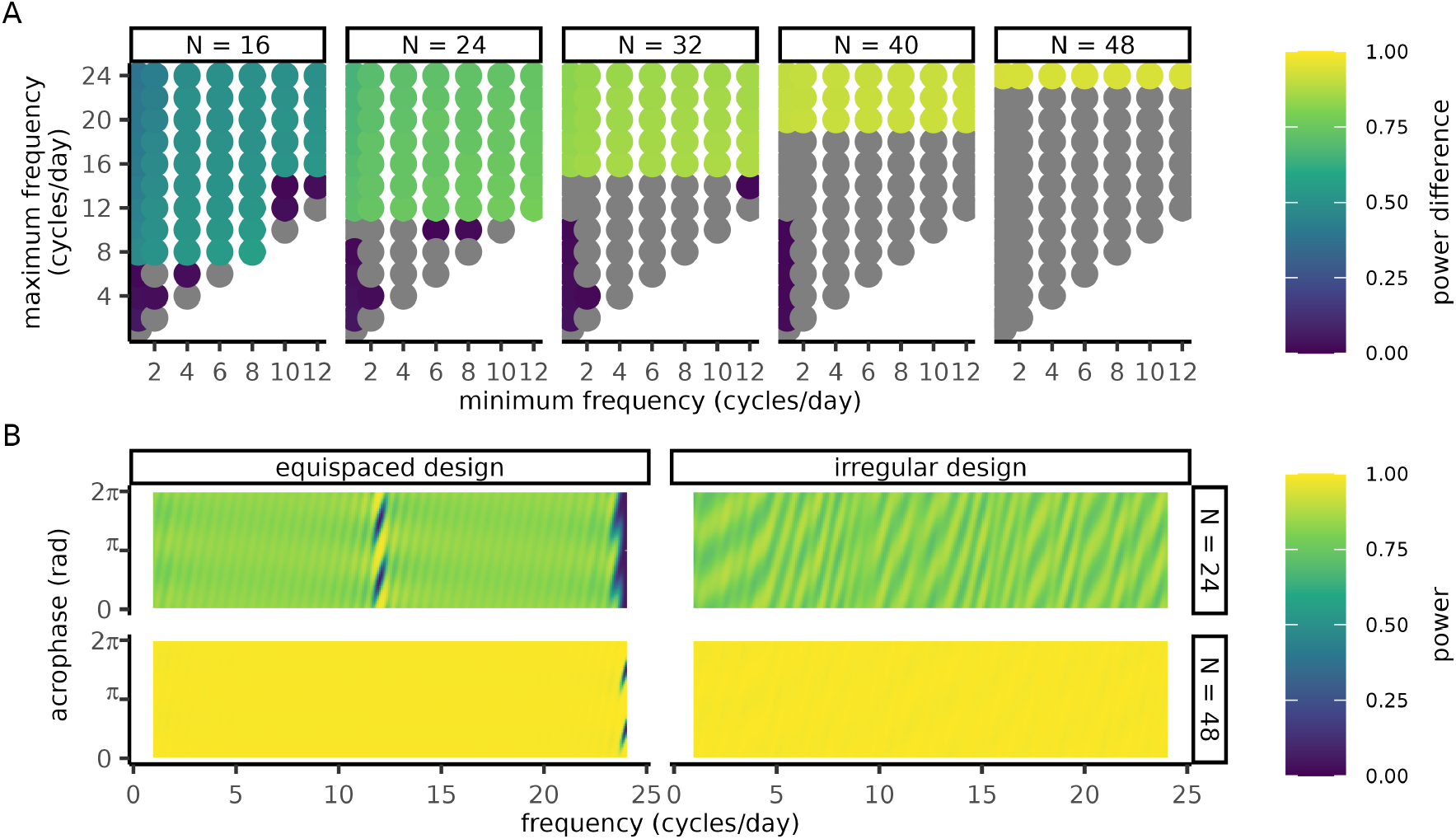
Irregular sampling improves power in experiments with period uncertainty. Irregular designs were optimized by differential evolution to detect signals whose frequency is in a window *f*_min_ ≤ *f* ≤ *f*_max_. **(A)** Differences in power between equispaced designs and optimized irregular designs where the *x* and *y* axes represent the upper (*f*_max_) and lower (*f*_min_) limits of each frequency range of interest. The color of each circle represents the difference in worst-case power between an irregular and equispaced design with the same number of measurements (panels). Grey circles correspond to negative power differentials. **(B)** Heatmap of power as a function of frequency (*x*-axis) and acrophase (*y*-axis) where color represents the power. **(A-B)** Simulated cosinor parameters: amplitude *A* = 1 and noise strength *σ* = 1. Each differential evolution was run with 1hr of compute time with parameters CR = 5 ×10^−2^, *N*_pop_ = 10^3^, *ε* = 5 ×10^−2^ (see Sec. 5.1). Details on power calculation at the Nyquist rate are given in Supp. Sec. S1.4.

In this article, we present a method for constructing experimental designs that are optimal for detecting unknown rhythms. We obtain an efficient way to generate such designs by deriving a closed-form expression for the statistical power of the cosinor model in Section 3.1 and focusing on designs where the least detectable signal of interest has the highest possible statistical power. Using this formula, we derived a condition for when equispaced designs fail to be optimal for rhythm discovery and investigated its consequences numerically in Sections 3.2-3.4. In Section 3.2 we present optimized designs for rhythm discovery across a range of candidate periodicities while in Section 3.3, we optimize for experiments where the period is known but the measurement time windows are restricted. In Section 3.4 we consider studies with strong prior knowledge of the system, so that only a discrete list of periods is of interest and demonstrates that certain periods can be measured together simultaneously without trade-offs in power. Our analysis and optimization methods generate more efficient experimental designs for rhythm discovery, and the computational methods can be applied to additional experimental contexts using the PowerCHORD (*P*ower analysis and *C*osinor design optimization for *HO*moscedastic *R*hythm *D*etection) open source code repository.

## 2 Background

Harmonic regression is a popular statistical framework for studying systems with oscillatory features [22]. We perform harmonic regression using the single-frequency cosinor model. For a MESOR (*M*idline *E*stimating *S*tatistic of *R*hythm) *Y*_0_, amplitude *A*, acrophase *ϕ*, and frequency *f*, the cosinor model takes the form

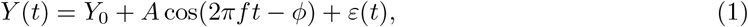

in which *ε*(*t*) ∼ 𝒩 (0, *σ*) is homoscedastic Gaussian white noise. Assuming the frequency *f* is fixed, Eq. (1) can be rewritten as

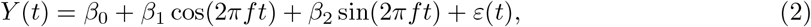

so that all unknown parameters appear linearly. Given data 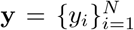 measured at times 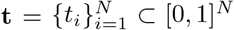, where *N* is the sample size of the experiment, the optimal least squares solution 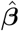 for estimating the coefficients ***β*** = (*β*_0_, *β*_1_, *β*_2_) appearing in Eq. (2) is

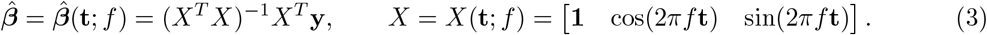

We assume **t** contains sufficiently many distinct measurement times for *X* to be of full column rank and Eq. (3) to be well-defined. For the cosinor model, the rank condition can be rephrased in terms of distinct phases of the cycle at which measurements are collected (Supp. Lemma S1.1). Estimates of the amplitude and acrophase of the signal can be computed from 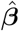,to obtain

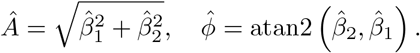

Using the cosinor model, rhythm detection can be formulated as a hypothesis test with null hypothesis *β*_1_ = 0 = *β*_2_ and alternative *β*_1_≠ 0 or *β*_2_ ≠0. The statistical power quantifies h ow reliably oscillations are detected by the experimental design and hypothesis test.

### Definition 2.1

(Statistical power of the cosinor model). Given a parametric model with parameters ***β*** ∈ℝ^*p*^, data *Y*∈ ℝ^*N*^ measured at times **t** ∈ℝ^*N*^, and a rejection region *R* ⊂ℝ^*N*^, the power of the hypothesis test is given by

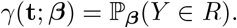

For a cosinor-based hypothesis test, the rejection region is 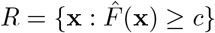,in which 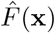 is the F-statistic

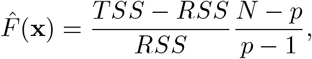

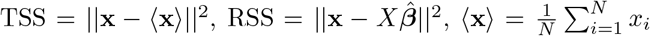,and *X* = *X*(**t**; *f*) is the cosinor design matrix and 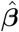 is the least-squares estimate of the parameters from Eq. (3).

Since the exact parameters of the signal are rarely well-known when the design is constructed, we seek designs that achieve high power across a range of parameter values. To this end, we quantify performance using the worst-case power of the design, meaning the lowest power across all signals of interest. Prioritization of worst-case rather than average power ensures that the power is above a known threshold for all relevant signals. A formal definition of worst-case power is given below. For the remainder of the paper we will simply use the terminology “optimal power designs” to refer to optimality with respect to a worst-case scenario.

### Definition 2.2

(Optimal worst-case power). Given a domain ℬ⊂ ℝ^*p*^ in parameter space and a design matrix *X* ∈ ℝ^*N ×p*^, an experimental design **t**^*^ achieves optimal worst-case power with respect to B if it satisfies

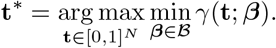

## 3 Results

### 3.1 Calculation and optimization of cosinor power

Power analysis is typically approached using Monte Carlo techniques. These techniques are transferable to power optimization, but introduce a significant computational burden. We took a more efficient route by deriving a closed-form expression for statistical power, stated in Theorem 3.1. The power expression is fast to evaluate, generalizes an earlier formula [23] which is only valid when measurements are equispaced (Supp. Fig. 2), and provides insight into the problem’s mathematical structure. The reader is directed to Supp. Sec. S1 for proofs of this result and all subsequent theorems.

**Figure 2:**
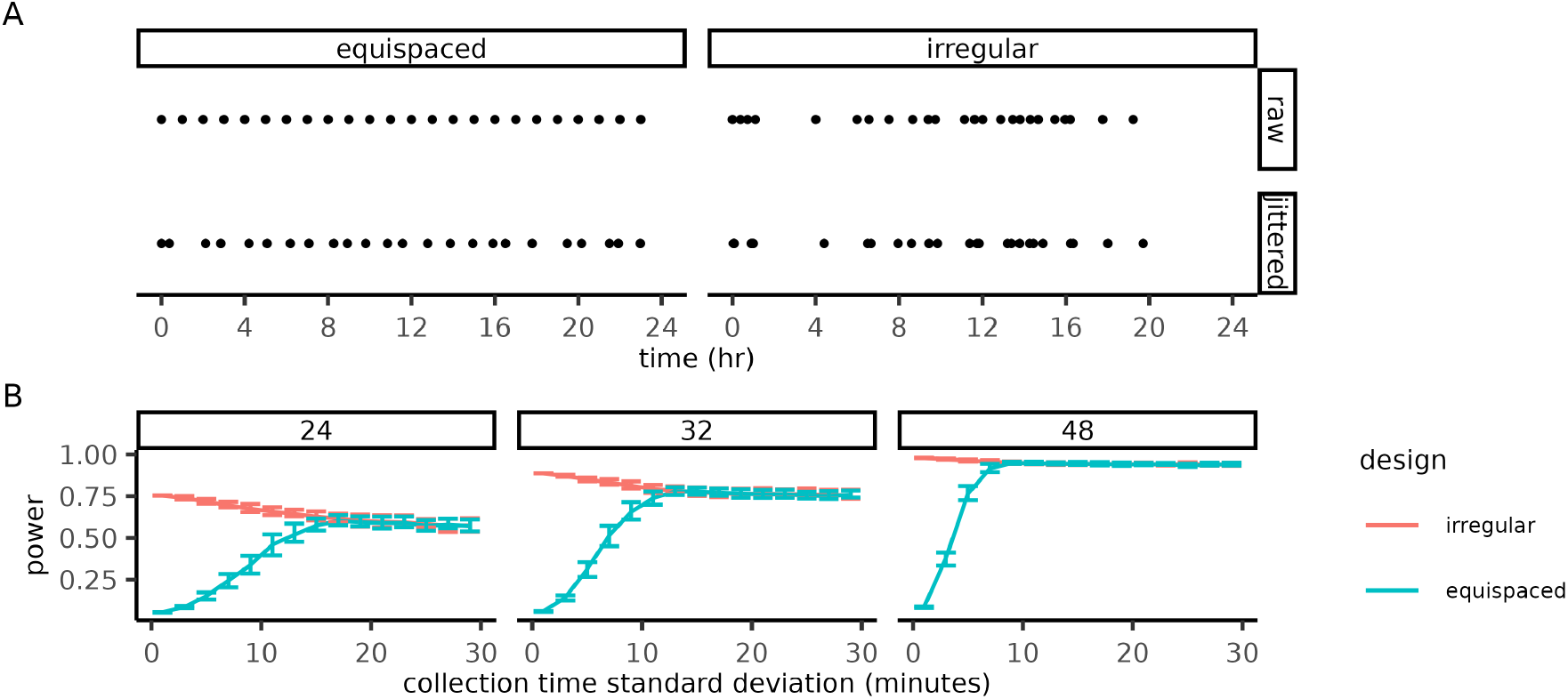
Irregular designs are robust to perturbations in measurement timing. **(A)** Measurement schedules before (top) and after (bottom) jittering. Jittered designs were generated by randomly perturbing the measurement times of equispaced and irregular designs with Gaussian white noise. **(B)** Worst-case power of jittered designs (x-axis) as a function of noise intensity (y-axis) for various sample sizes (panels). The bars indicate the interquartile range for ensembles (*n* = 100) jittered designs. Irregular designs were optimized for the frequency window *f*_min_ = 1 and *f*_max_ = 24 using differential evolution with same parameters as Fig. 1. Power was calculated assuming an amplitude *A* = 1.

#### Theorem 3.1

(Power of the one-frequency cosinor model). *Consider the one-frequency cosinor model*

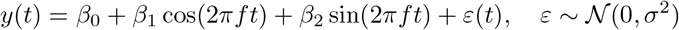

*applied to data* 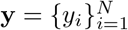 *collected at distinct times* 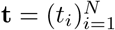 *with N >* 3. *Suppose the following hypotheses are tested using an F-test*

- *null hypothesis H*_0_ : *β*_1_ = 0 = *β*_2_,
- *alternative hypothesis H*_1_ : *β*_1_ ≠ 0 *or β*_2_ ≠ 0.

*Given parameters* ***β*** = (*β*_0_, *β*_1_, *β*_2_), *the power γ of this hypothesis test is given by*

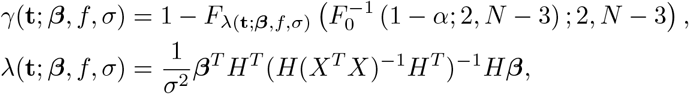

*in which X* = *X*(**t**; *f*) = 1 cos(2*πf* **t**) sin(2*πf* **t** *is the design matrix, α* ∈ (0, 1) *is the type I error rate, F*_*λ*_(*x*; *n*_1_, *n*_2_) *is the noncentral F-distribution with* (*n*_1_, *n*_2_) *degrees of freedom and noncentrality parameter λ, F*_0_(*x*; *n*_1_, *n*_2_) = *F* (*x*; *n*_1_, *n*_2_) *is the F-distribution, and*

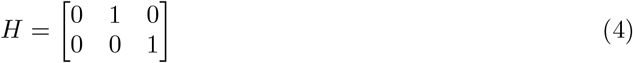

*is the hypothesis matrix*.

Using Theorem 3.1, power optimization can be equivalently formulated as an eigenvalue programming problem. The resemblance of this eigenvalue problem to the Elfving optimality criterion from optimal experimental design [20] leads to a simple condition for the optimality of equispaced designs. When the optimality condition holds, the power of equispaced designs is also guaranteed to be independent of the signal’s acrophase (Supp. Fig. 3A). The eigenvalue problem and optimality condition are stated in Theorem 3.2. If an experiment cannot satisfy the optimality condition, optimal designs can be constructed numerically using Corollary 3.2.1.

**Figure 3:**
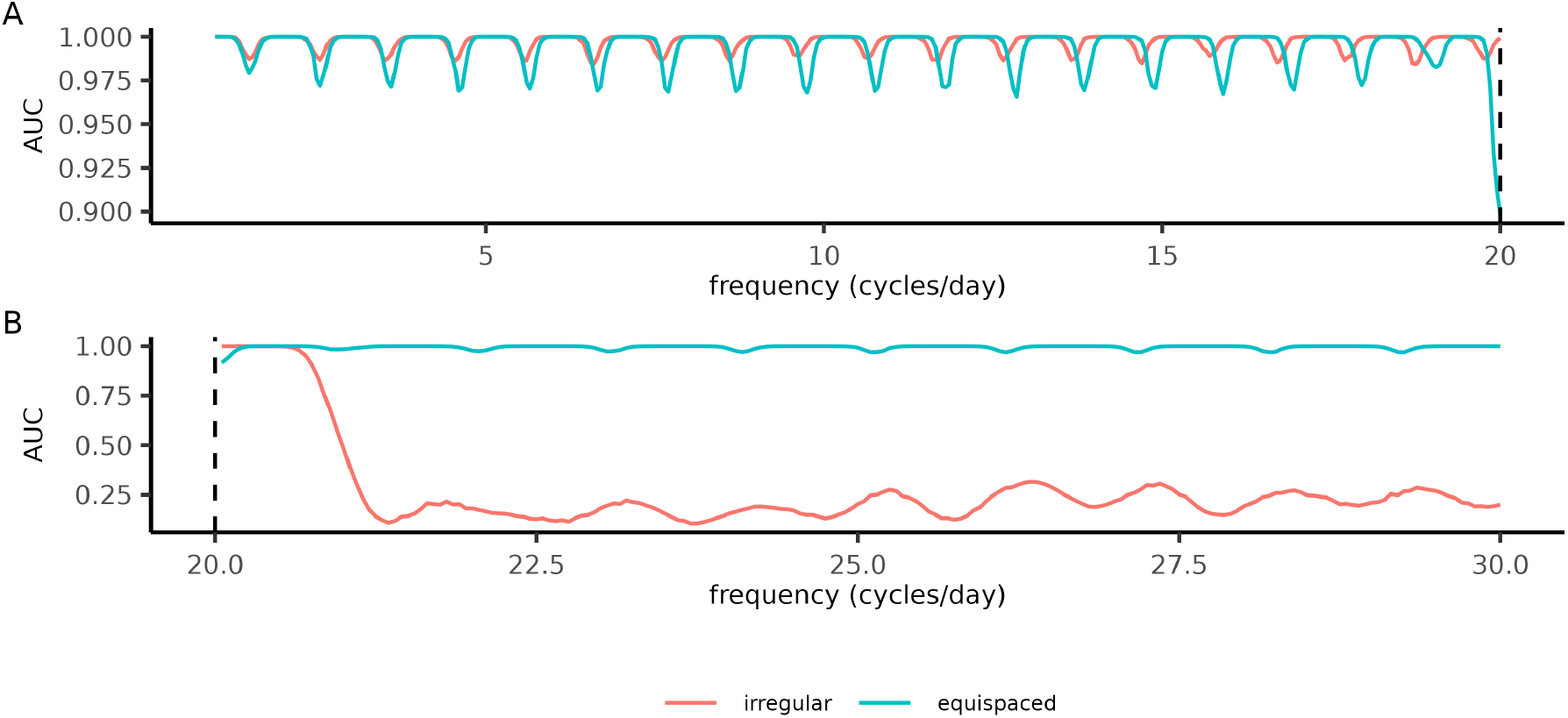
Irregular designs improve simulated periodogram analysis at frequencies up to the Nyquist rate. **(A)-(B)** Oscillations were detected using periodogram analysis with measurements (*N* = 40 samples) from either an equispaced or irregular design optimized for the frequencies 1≤ *f* ≤20. Each signal in the dataset was assigned an oscillatory (amplitude *A* = 2, noise strength *σ* = 1) or non-oscillatory (amplitude *A* = 0, noise strength *σ* = 1) state with equal probability. The acrophase of the oscillatory signals was assigned uniformly at random (*ϕ* ∼Unif([0, 2*π*))). **(A)** Performance of the irregular and equispaced designs at detecting oscillations across frequencies (x-axis) included in the optimization, summarized by the AUC score (y-axis) of a receiver operator characteristic curve with p-values generated from a Lomb-Scargle periodogram. The AUC score for each frequency was computed by testing for oscillations in a dataset of oscillatory and white-noise signals (*n* = 10^4^ signals per dataset). The dashed line indicates the Nyquist rate of the equispaced design. **(B)** The same analysis as (A) but at frequencies above the Nyquist rate of the equispaced design. Periodogram analysis was performed using the lomb library [28] and AUC scores were computed using the `pROC` library [29]. Irregular designs were generated using the same differential evolution parameters as in Fig. 1.

#### Theorem 3.2

(Optimality condition for equispaced designs). *Suppose a study aims to detect signals of a specific frequency and unknown acrophase. Maximizing the worst-case power across all acrophases is equivalent to the following eigenvalue optimization problem*

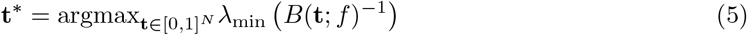

*in which*

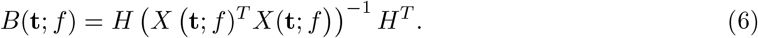

*If it is possible to collect N >* 3 *equispaced measurements per cycle, then equispaced designs maximize Eq*. (5). *Moreover, the power of such designs is constant as a function of acrophase with power corresponding to a noncentrality parameter*

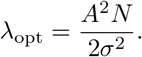

#### Corollary 3.2.1.

*If the frequency is assumed to lie in a specific window f* ∈ [*f*_min_, *f*_max_] *and the acrophase is unknown, then maximizing the worst-case power is equivalent to the following eigenvalue programming problem*

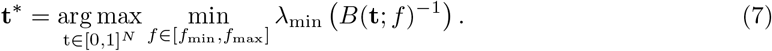

*The value achieved by this optimal solution* **t**^*^ *is bounded by*

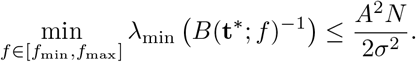

PowerCHORD generates designs by maximizing Eq. (7) using a brute-force search, differential evolution, or mixed-integer conic programming. Brute-force searches are computationally tractable if the sample size is low and the measurements are confined to an underlying grid (e.g. only sampling on the hour). For higher sample sizes, differential evolution provides an estimate of the power improvement and mixed-integer conic programming provides more refined information and better structured solutions. The three methods work directly with the eigenvalue problem and their results can be converted back into power using the following identity

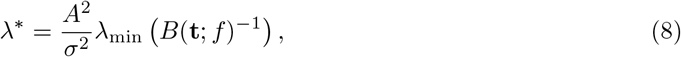

in which *λ*_min_ (*B*(**t**; *f*)^−1^) is the minimal eigenvalue from Eq. (5) and *λ*^*^ is the lowest value of the noncentrality parameter across all acrophases for signals of amplitude *A* and frequency *f*. Justification for Eq. (8) is given in the proof of Supp. Lemma S1.12. In our analysis, we set *A* = *σ* so that we may refer to the noncentrality parameter and minimum eigenvalue *λ*_min_ (*B*(**t**; *f*)^−1^) interchangeably. We also make use of the fact that the noncentral F distribution is a monotone function of its noncentrality parameter (Supp. Sec. S1.2) and therefore refer to the power of a design and its noncentrality parameter interchangeably.

### 3.2 Irregular designs improve power across frequency windows

We assume the study seeks to achieve high power across a frequency window *f*_min_ ≤ *f* ≤ *f*_max_. Using PowerCHORD’s differential evolution method, we constructed irregular designs for various frequency windows and sample sizes, which improved power for most cases (Fig. 1A). The greatest improvements to power were observed when the frequency window included the Nyquist rate of the equispaced design. Optimization of other windows did not improve upon equispaced designs or only produced a slight change in the power. In comparison to equispaced designs, the power of irregular designs had much weaker power fluctuations around the Nyquist rate (Fig. 1B; Supp. Fig. 3B). Away from the Nyquist rate, the equispaced and irregular designs exhibited power fluctuations that diminish in intensity as the sample size increases. Hence, for a large enough sample size, irregular designs improve worst-case power while maintaining homogeneous power levels throughout the frequency-acrophase plane.

Some of the measurements in irregular designs are spaced farther apart than in equispaced designs of the same sample size. These measurement gaps could make the design undesirably sensitive to the timing of specific measurements. To investigate this sensitivity, we generated “jittered” irregular and equispaced designs by adding Gaussian white noise to each measurement time (Fig. 2A) and calculated the worst-case power of the increasingly perturbed designs. As the noise intensity increased, the power improvements of the irregular design declined as both designs converged in power as measurement times were effectively being sampled from the same distribution (Fig. 2B). For a sample size of 24, the power difference between the irregular and equispaced designs reached half of its full difference when the measurement error was 9 minutes, while for a larger sample size of 48 this difference diminishes more quickly at a measurement error of 5 minutes. Irregular designs always outperformed the eventual random design (Δ = 0.18, 0.14, 0.04 for *N* = 24, 32, 48, respectively). Since timing error is likely to be on the order of minutes, our power improvements are robust to errors that occur on a realistic timescale.

Our irregular designs were optimized for cosinor analysis, yet the data they produce may be used for other purposes such as spectral analysis. To determine the applicability of our irregular designs in such a context, we performed periodogram analysis on simulated datasets. We made use of the Lomb-Scargle periodogram, a common method developed for the study of irregularly sampled data [24,25], and applied a hypothesis test derived from this method [26]. We generated independent simulated datasets for a range of frequencies, each made up of white noise and oscillatory signals with uniformly random acrophases. We quantified performance using the area under an ROC (receiver operator characteristic) curve [27] because it can be calculated without knowing the false positive rate, and this rate could be affected by switching from cosinor to periodogram analysis. Relative to equispaced designs, irregular designs exhibited weaker fluctuations in their AUC (area under the curve) score at frequencies below the Nyquist rate (Fig. 3A). At the Nyquist rate, the irregular designs dramatically outperformed the equispaced design. Irregular designs performed well only at the frequencies included in their optimization (Fig. 3B), emphasizing the importance of choosing a realistic frequency range before optimizing an experimental design. Although they were constructed for cosinor-based analysis, PowerCHORD designs remain applicable in a broader statistical context.

### 3.3 PowerCHORD recovers power lost to timing constraints

Equispaced designs can provide optimal power in fixed-period experiments, but they may not always be the most convenient option. For instance, a circadian study with human participants may aim to minimize the number of times a participant is awoken during the night for sample collection. Since strong acute disruptions of sleep have been shown to impact various biological measurements [30,31], accurate characterization of naturally occurring oscillations should avoid disruptions to a 6-12hr window of a subject’s regular sleep routine. Incorporating a rest-window reduces the power relative to an equispaced design, so we aimed to determine how much power could be recovered by optimizing the measurement times under the timing constraint.

To simplify the design space, we assumed that measurements could only be collected at times aligned with a half-hour grid. Consequently, it was possible to find optimal designs by running a brute-force search (Sec. 5.2). To gauge the benefit of timing optimization, we compared the optimal designs to “naive” designs in which all measurements are equispaced outside the rest-window (Fig. 4A). Relative to the naive designs, the power of the constrained-optimal designs was considerably less sensitive to the acrophase of the signal (Fig. 4B). This difference in sensitivity can be observed in the peak-to-trough power variability of the two designs (Δ_naive_ = 22.93%, Δ_optimal_ = 0.35%; 8hr rest-window, amplitude *A* = 2.5). To estimate how much the constraints limit the power, we compared the noncentrality parameters of the constrained-optimal and naive designs to an equispaced design. Negligible power was lost in the constrained-optimal design when imposing the shortest window duration. As the window length increased, both the constrained-optimal and naive designs lost power (Fig. 4C). Still, the constrained-optimal designs out-performed the naive designs across the windows under consideration, suggesting that timing optimization can help studies accommodate timing constraints without unnecessarily reducing their power.

**Figure 4:**
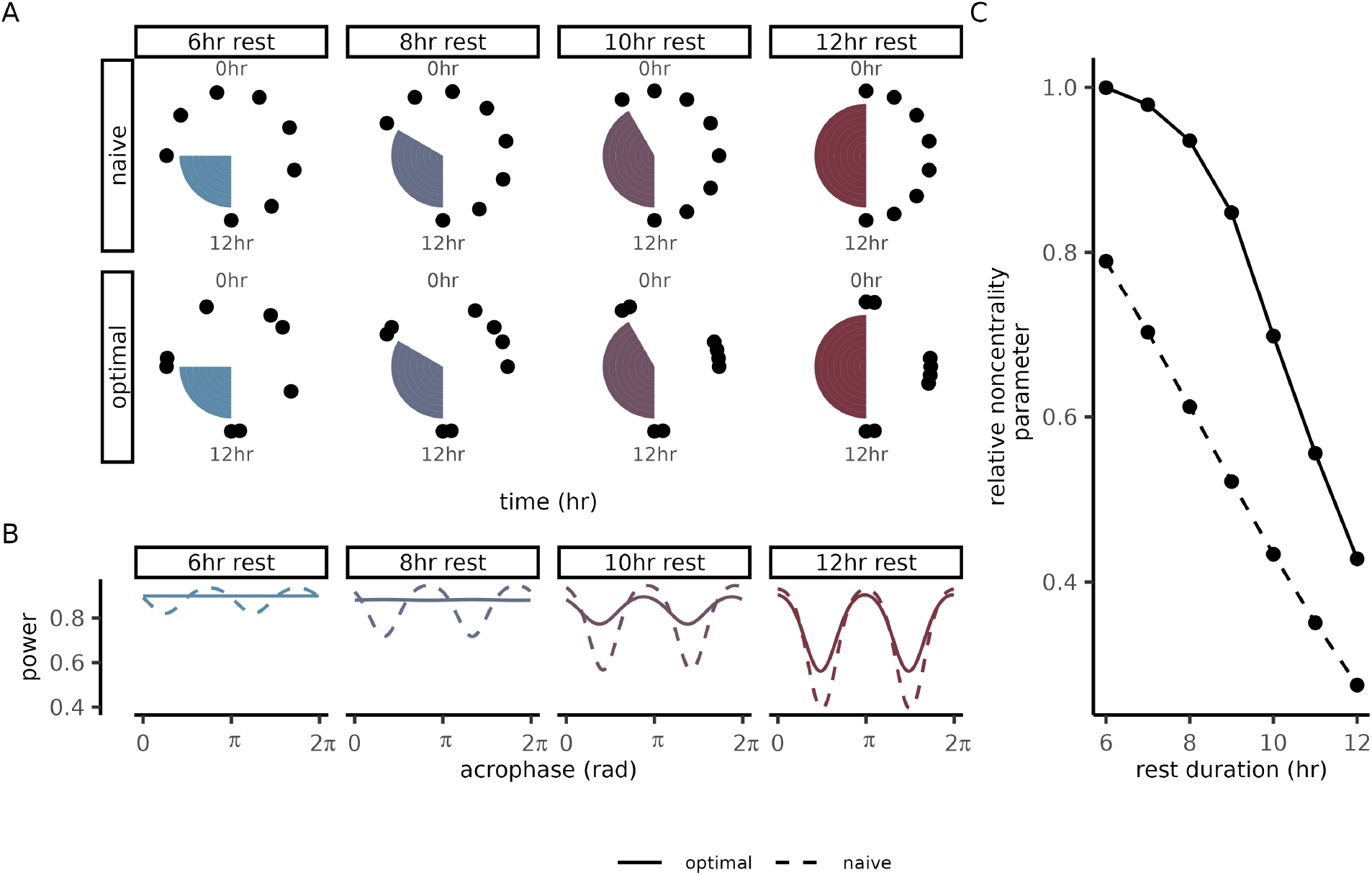
Circadian studies can balance power with rest-window duration using timing optimization. **(A)** Measurement times for optimal and naive designs plotted as phases of a 24hr cycle. Each panel corresponds to a different duration of the rest-window (shaded region), during which no samples can be collected. Optimal designs were generated using a brute-force search with *N*_*t*_ = 48 points in the temporal discretization (Sec. 5.2). Naive designs were constructed by distributing all measurements outside the rest window at equal time-intervals. **(B)** Power (y-axis) as a function of acrophase (x-axis) for various durations of the rest-window. The color of each curve represents rest-window duration and the line-type represents if the design is optimal or naive. **(C)** The noncentrality parameter (y-axis) as a function of rest-window duration (x-axis) for naive and optimal designs. Noncentrality parameters are reported relative to the noncentrality parameter of an unconstrained equispaced design with the same measurement budget (*N* = 8 samples).

### 3.4 Globally optimal designs for discrete period uncertainty

Brute-force searches quickly become intractable at high sample sizes (Supp. Fig. 4). Our third method is a mixed-integer conic program (Theorem 3.3) that continues to be informative about global optimality and remains applicable at higher sample sizes. After simplifying the design problem to only include a small number of frequencies (i.e. discrete period uncertainty), the conic program converged and provided numerical evidence of global optimality. In contrast to the previous two optimization methods, which relied only on the computational efficiency of our power formula, the construction of the conic program relies on the mathematical structure present in the formula (Sec. 5.3).

#### Theorem 3.3

(Mixed-integer conic program for power optimization). *Worst-case power maximization with discrete time*

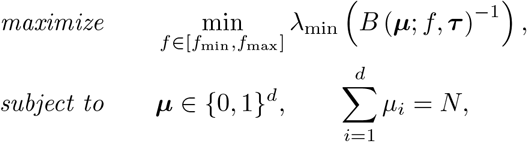

*is equivalent to the following mixed-integer conic programming problem*

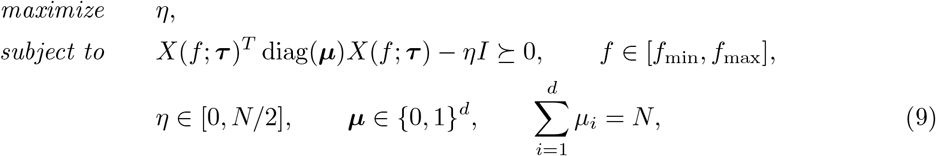

*in which X*(*f*, ***τ***) *is the design matrix of the one-frequency cosinor model evaluated at frequency f on the partition* ***τ***, *and B* (***µ***; *f*, ***τ***) *is given by*

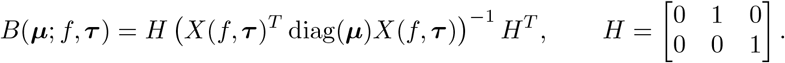

Optimal designs were generated by maximizing Eq. (9) using the CUTSDP method in YALMIP [32] together with Gurobi as a backend solver [33]. The widest frequency separation corresponded to resolving a pair of periods *T* = 24hr and *T* = 2hr with a sample size of *N* = 12 measurements. Interestingly, the optimal solution contained repeating patterns in its measurement schedule (Fig. 5A; Supp. Fig. 5A). While the equispaced design suffered from a low noncentrality parameter at the 2hr rhythm (Fig. 5B), the optimal design reached its theoretical maximum value (Theorem 3.2) at both frequencies of interest. This ability to resolve both frequencies without sacrificing power at either frequency was not observed in 10,000 randomly generated designs (Supp. Fig. 6A) or when additional frequencies were included in the design problem (Supp. Fig. 6B). Closer examination of the bifrequency optimal designs revealed a simple explanation for their ideal power balance; they can be split into equispaced designs when visualized as phases of each period in the design problem (Fig. 5C-D) and therefore inherit the optimality properties of equispaced designs (Supp. Sec. S1.5). Since optimal designs can be constructed by simply ensuring that they satisfy this equiphase property, we used this approach to construct optimal designs for detecting circadian, circalunar, and circannual rhythms in a 12 month experiment (Fig. 5E-F; Supp. Fig. 6C).

**Figure 5:**
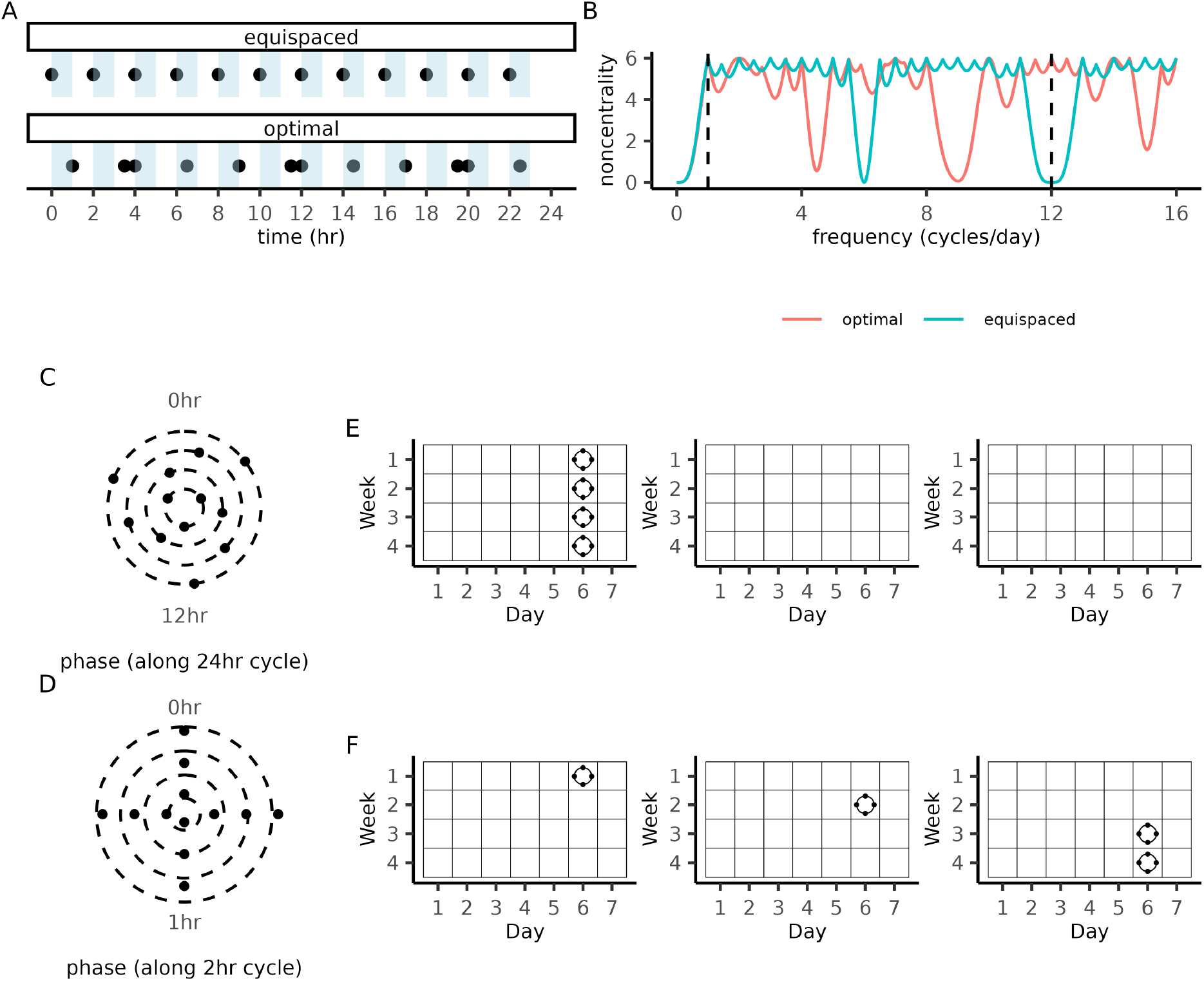
Globally optimal designs for discrete period uncertainty. **(A)** Measurement collection times of an equispaced design (top) and a bifrequency optimal design (bottom) for detecting 2hr and 24hr rhythms. **(B)** Power (y-axis) as a function of frequency (x-axis) for the equispaced and bifrequency optimal design. The two frequencies included in the optimization problem are indicated by the dashed vertical lines. **(C)** Bifrequency optimal design plotted as phases of a 24hr cycle. The values of the radial coordinate were chosen to emphasize that the design can be split into four equiphase designs, each containing three measurement times. **(D)** Bifrequency optimal design plotted as phases of the 2hr cycle. The radial position of each point represents its phase along the 24hr cycle. **(E)** Days marked with circles indicate that *n* = 4 equispaced circadian measurements are to be collected. Repeating this schedule four times over the course of a year, produces a trifrequency optimal design with measurements equispaced along circadian (24 hr), circalunar (28 day), and circannual (12 × 28 = 336 day) cycles. (**F)** An alternative measurement schedule that distributes the measurements across all three months while maintaining the equiphase property.

## 4 Discussion

We presented a closed-form expression for the statistical power of cosinor based rhythm detection which holds for arbitrary measurement times. This formula enables exploration of the rhythm detection design space in a manner that was infeasible with Monte-Carlo power estimation. This formula will be useful for regular statistical practice to answer questions such as the phase-detection bias introduced by the loss of a sample, or in datasets which may not have been optimally designed (e.g. a postmortem dataset [34]). Using our closed-form power expression, we proved that any given periodicity can be optimally detected by an equispaced design with *N >* 3 measurements per cycle. This optimality condition implies that certain groups of periods can be investigated together without any trade-off in power. In particular, the power will be phase-independent provided that the measurements decompose into equispaced designs when viewed as phases of each period under consideration.

Application of PowerCHORD methods to various experimental scenarios improved the power relative to equispaced designs. When designs were optimized for a continuous range of periods, the improved power was robust to small changes in measurement timing and robust to the application of periodogram rather than cosinor analysis. Under the simplifying assumption that measurements are confined to an underlying grid, two of our three optimization methods (brute-force search and conic programming) produced globally optimal designs. The optimal designs produced by the brute-force searches exhibited a minimal loss in power relative to equispaced while removing the need for costly and logistically difficult overnight visits and avoiding sleep disruption biases. In the conic programming case, the globally optimal designs had noncentrality parameter *λ* = *N/*2 at all frequencies of interest and hence their performance was not limited by the restriction to an underlying grid. Examination of the bifrequency optimal designs revealed an “equiphase” property and led to simple optimal designs for simultaneous investigation of circadian, circalunar, and circannual rhythms.

It may be possible to generate optimal designs in more challenging experimental contexts by improving the optimization methods in PowerCHORD. The brute-force searches could be accelerated to run at larger sample sizes using more efficient algorithms for parameterising the design space [35] and related methods in optimal experimental design [36–38] could be adapted to our problem. While our analysis was focused on how measurement timing influences worst-case power, there are many closely related questions that could be considered in future work. First, the number of biological and technical replicates could be treated as decision variables in the optimization problem [14, 39] to further improve power. Second, figures of merit relevant to the accuracy of parameter estimation or periodogram methods [26, 28, 40, 41] could be prioritized in the study design. Third, timing optimization could be applied to adaptive study design. In this setting, the frequency prior would be updated using Bayesian inference and influence the optimization of measurement times for the next experiment [42]. Beyond the scope of optimization, further improvements to rhythm detection can be achieved by higher quality data, innovative analytical methods and leveraging prior knowledge of the system [43].

In summary, we have demonstrated the benefits of integrating power optimization in the design of rhythm discovery experiments. Our methods are broadly applicable, and can be tailored to specific experimental applications using our open-source PowerCHORD repository. Such studies will be more efficient than traditional approaches for exploring ranges of periods, and may expedite the discovery of novel biological rhythms.

## 5 Methods

### 5.1 Differential evolution

Differential evolution algorithms initialize a population of candidate solutions at random. Successive generations are constructed by taking component-wise linear combinations of the parents according to an algorithm-specific update rule. While many variations on this core idea have been studied [44], we chose to implement a relatively simple version of the algorithm.

Given an objective function *J* : ℝ^*N*^ → ℝ, our implementation of differential evolution requires four hyper-parameters: the population size *N*_pop_, the differential weight *ε* ∈ [0, 1], the crossover probability *CR* ∈ [0, 1], and the number of iterations *N*_iter_. At initialization, the population is represented by a matrix 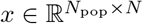,in which *N* is the sample size, whose entries are independently and identically distributed as *x*_*ij*_ ∼ unif(0, 1). To produce the next generation, the state of the *i*-th member of the population is updated by generating a random vector *u* ∼ unif(0, 1), sampling three indices *a, b, c* ∈ {1, …, *N*_pop_} without replacement to obtain

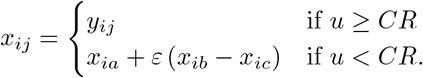

If *J*(*y*_*i*_) *< J*(*x*_*i*_), then the new state is accepted. The system repeats this process until a total of *N*_iter_ generations have been produced. The highest scoring individual of the terminal population is then returned. A summary of the algorithm in pseudo-code is given below.

#### Algorithm 1

Differential evolution in PowerCHORD

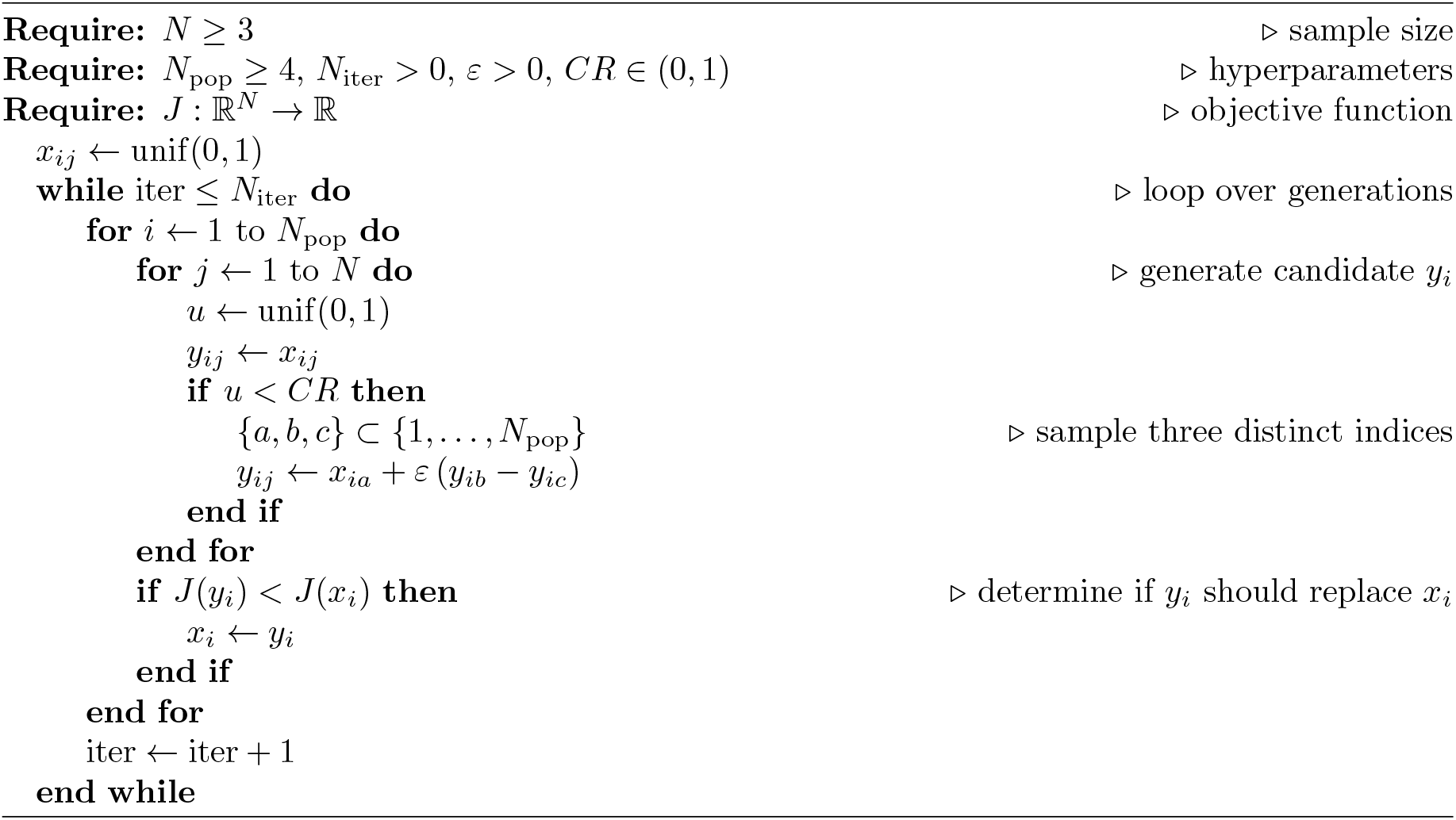

### 5.2 Brute-force search

Suppose measurements can only be collected at times belonging to a fixed partition

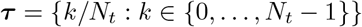

in which *N*_*t*_ is the coarseness of the partition. With measurement times confined to a fixed grid, designs are naturally represented by binary vectors 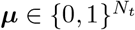 which satisfy ∑_*i*_ *µ*_*i*_ = *N*. Provided that *N* and *N*_*t*_ are not too large, it is feasible to search the entire space of binary vectors. Since any design 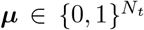 will achieve equal performance to all of its cyclic translations, we need only consider equivalence classes of such binary vectors up to rotational symmetry. Following the convention from combinatorics, these equivalence classes are referred to as fixed-density binary necklaces. For a partition of coarseness *N*_*t*_ and sample size *N*, the number of equivalence classes of such necklaces is given by a sum over the factors of the greatest common divisor of the sample size and the partition coarseness

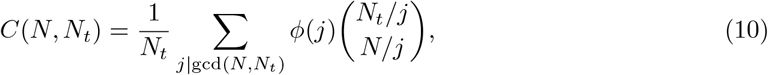

in which *ϕ*(·) is Euler’s totient function [45]. We recommend restricting *N*_*t*_ ≤ 72 and *N* ≤ 10 since this function grows rapidly with *N* and *N*_*t*_ (Supp Fig. 4). An efficient algorithm for generating a representative from each equivalence class is given in [45]. We use their C implementation to generate the designs and search the design database to identify representatives of equivalence classes of optimal solutions.

There is an additional reflection symmetry present in the problem which could further improve the performance of the brute-force search. Notice that the time-reversal of any design will still have equivalent power, hence designs could be considered equivalent up to rotational and reflectional symmetry. In the terminology of combinatorics, these larger equivalence classes are known as “fixed-density binary bracelets”. Efficient algorithms for generating representatives of the bracelets have been proposed [35] and would improve the performance of PowerCHORD if they were included.

### 5.3 Mixed-integer conic programming

We provide a proof of Theorem 3.3 which justifies our use of mixed-integer conic programming. We simplify the optimization problem using the following lemma.

#### Lemma 5.1

(Schur positivity, [46, Section A.5.5]). *Let Y* ∈ ℝ^*n×n*^ *be a symmetric matrix partitioned as*

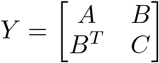

*with* det *A*≠0 *and let S* = *C* − *BA*^−1^*B*^*T*^ *be the Schur complement of Y*. *Then we have Y* ≻ 0 *if and only if A* ≻ 0 *and S* ≻ 0.

**Proof of Theorem 3.3**. Let *X* = *X*(***τ*** ; *f*) and 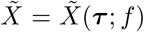.We seek a binary vector ***µ***^*^ that satisfies

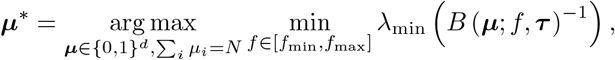

in which *B*(***µ***; *f*, ***τ***)^−1^ can be expressed as

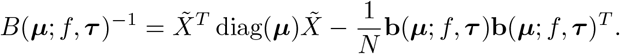

with the latter equality justified by Lemma S1.13. Restricting frequency to a discrete set of frequencies of interest {*f*_1_, …, *f*_*M*_}, we arrive at an optimization problem of the form

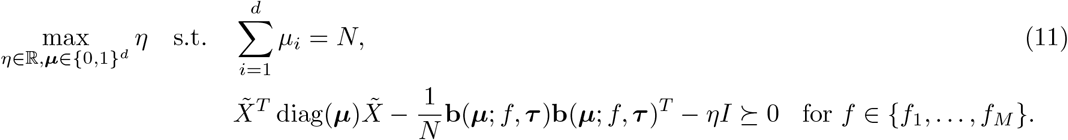

The quadratic dependence on ***µ*** in Eq. (11) can be reduced to a linear dependence by applying Lemma 5.1 with the matrix *Y* given by

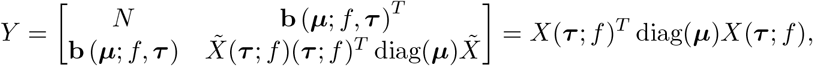

which gives

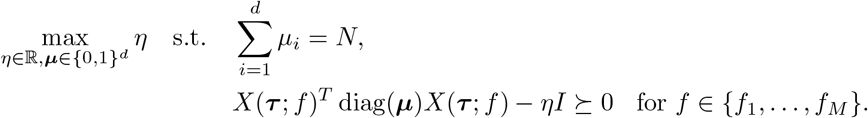

## 6 Code Availability

Implementations of PowerCHORD in R and MATLAB are available through the first author’s GitHub. The power analysis features are included in both the R and MATLAB implementations. Only the MATLAB implementation includes GPU acceleration and provides access to the external YALMIP library necessary for mixed-integer conic programming.

## 7 Acknowledgements

TS is supported by an NSERC Canada Graduate Scholarship. This work was supported by the Krembil Foundation, Toronto, Canada, the Future Biomedicine Charity Fund, Vilnius, Lithuania, and by Research Council of Lithuania under the Programme “University Excellence Initiatives” of the Ministry of Education, Science and Sports of the Republic of Lithuania (No. 12-001-01-01-01 “Improving the Research and Study Environment”; project No S-A-UEI-23-10). AP is a Marius Jakulis Jason Foundation scholar. ARS acknowledges the support of the Natural Sciences and Engineering Research Council of Canada (NSERC): RGPIN-2019-06946. This research was enabled in part by support provided by Compute Ontario and the Digital Research Alliance of Canada. We give thanks to Michael Tackenberg and John Limanto for providing thorough and insightful feedback on this manuscript.

## Supplementary material

### S1 Proofs of main results

#### S1.1 Closed-form power expression

The power formula in Theorem 3.1 follows from computing the distribution of the F-statistic under the null and alternative models. We show these computations in detail since self-contained proofs are unavailable in the literature. The first three lemmas are concerned with elementary properties of the design matrix *X* and the hypothesis matrix *H*.

##### Lemma S1.1.

*Let X* = *X*(**t**; *f*) ∈ ℝ^*N ×*3^ *be the design matrix of the cosinor model and suppose at least three components of the vector* **z** = *e*^2*πif***t**^ ∈ ℂ^*N*^ *are distinct, then X has full column rank*.

*Proof*. Without loss of generality, assume the first three components *z*_1_, *z*_2_, *z*_3_ of **z** ∈ ℂ^*N*^ are distinct. Let *θ*_1_, *θ*_2_, *θ*_3_ ∈ [0, 2*π*) be the corresponding angles *θ*_*j*_ = Arg(*z*_*j*_). To verify that *X* is of full column rank, it suffices to show that the submatrix

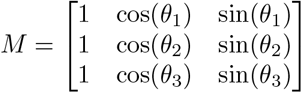

has nonzero determinant. Assume without loss of generality that *θ*_3_ = 0 and compute the determinant of *M* to obtain

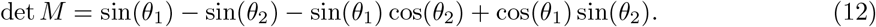

Suppose towards contradiction that det *M* = 0 and rearrange Eq. (12) to show

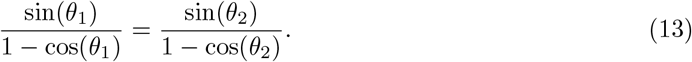

The restriction of the function 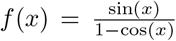 to *x* ∈ [0, 2*π*) is strictly monotone and therefore injective. Hence Eq. (13) implies *θ*_1_ = *θ*_2_, a contradiction.

##### Lemma S1.2.

*Let X* ∈ ℝ^*n×p*^ *with* rank(*X*) = *p* ≤ *n then X*^*T*^ *X is symmetric positive definite*.

*Proof*. Since *X*^*T*^ *X* is symmetric, there exists an orthonormal eigenbasis **v**_1_, …, **v**_*p*_ and real eigenvalues *λ*_1_, …, *λ*_*p*_ such that *X*^*T*^ *X***v**_*i*_ = *λ*_*i*_**v**_*i*_ for *i* = 1, …, *p*. Since rank(*X*) = *p*, we know *X***v**_*i*_ ≠ 0 and so

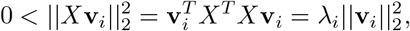

which forces *λ*_*i*_ *>* 0 for *i* = 1, …, *p*.

##### Lemma S1.3.

*Suppose A* ∈ ℝ^*p×p*^ *is symmetric positive definite and H* ∈ ℝ^*q×p*^ *with* rank(*H*) = *q. If q* ≤ *p then B* = *HAH*^*T*^ *is symmetric positive definite*.

*Proof*. Symmetry is apparent, so it suffices to show that **x**^*T*^ *HAH*^*T*^ **x** *>* 0 for any **x** ∈ ℝ^*q*^. Since rank(*H*) = *q*, we know *H*^*T*^ **x**≠0 for all nonzero **x** ∈ ℝ^*q*^. Positivity of *HAH*^*T*^ now follows from the positivity of *A*, since (**x**^*T*^ *H*)*A*(*H*^*T*^ **x**) *>* 0 for all **x** ∈ ℝ^*q*^.

The next lemma and its corollary will be useful for computing the null and alternative distributions of the F statistic. In particular, it is useful to recognise that the definition of the F-statistic most frequently applied in experiments can also be written in a form more suitable for mathematical proofs.

##### Lemma S1.4.

*Given a design matrix X* ∈ ℝ^*n×p*^ *with* rank(*X*) = *p and a hypothesis matrix H* ∈ ℝ^*q×p*^ *with q* ≤ *p* ≤ *n, define the following linear subspaces*

- *V*_*u*_ = {*X****β*** : ***β*** ∈ ℝ^*p*^} = Col(*X*),
- *V*_*r*_ = {*X****β*** : ***β*** ∈ ℝ^*p*^, *H****β*** = 0},
- *V*_*ℓ*_ = {*X****β*** : ***β*** ∈ ℝ^*p*^, *H****β*** 0}.

*Let P*_*u*_, *P*_*r*_, *and P*_*ℓ*_ *be the orthogonal projectors onto the respective subspaces. The projector P*_*ℓ*_ *satisfies*

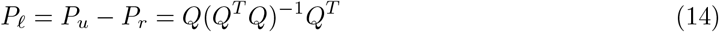

*in which Q* = *X*(*X*^*T*^ *X*)^−1^*H*^*T*^.

*Proof*. Since *V*_*u*_ can be expressed as a direct sum *V*_*u*_ = *V*_*ℓ*_ *⊕ V*_*r*_, we clearly have *P*_*ℓ*_ = *P*_*u*_ − *P*_*r*_. To verify the second equality, it suffices to show that the columns of *Q* span *V*_*ℓ*_. To confirm this, we verify the equivalent condition 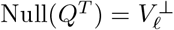.We may write 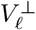 as 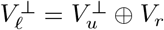.Any **y** ∈ *V*_*r*_ can be expressed as **y** = *X****β*** so *Q*^*T*^ **y** = *H****β*** = 0. Similarly 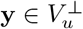 must satisfy *P*_*u*_**y** = *X*(*X*^*T*^ *X*)^−1^*X*^*T*^ **y** = 0 and hence *Q*^*T*^ **y** = 0 and we see 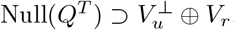. For the reverse containment, notice that any 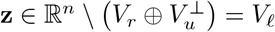,must satisfy **z** = *X****β*** and *H***z** ≠ 0. It follows that **z** *∉* Null(*Q*^*T*^) because

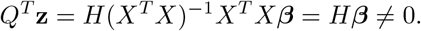

Since 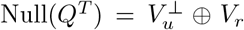,the rank-nullity theorem implies that Col(*Q*) = *V*_*ℓ*_ and the second equality of Eq. (14) follows.

##### Corollary S1.4.1.

*Let X* = *X*(**t**; *f*) *be the design matrix of the cosinor model. Given data* **y** ∈ ℝ^*n*^, *the total sum of squares*

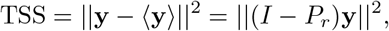

*and the residual sum of squares*

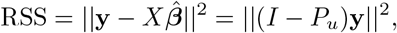

*in which* 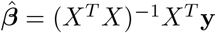 *is the least squares estimator and X is the one-frequency cosinor design matrix, the following three definitions o f the F -statistic are equivalent*

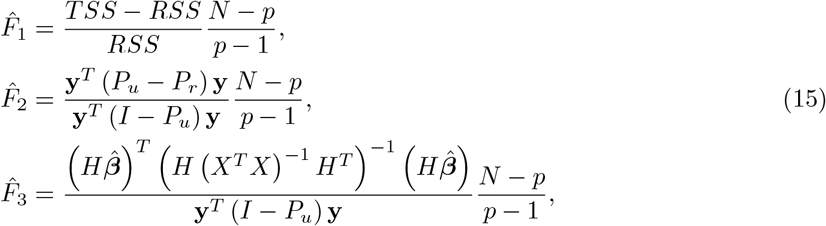

*in which* 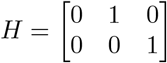 *is the rhythm-detection hypothesis matrix*.

The linear algebraic tools assembled so far allow us to deduce the null and alternative distributions of the F-statistic by simply applying Cochran’s theorem.

##### Theorem S1.5

(Cochran (1934), [47]). *If* **x** ∈ ℝ^*n*^ ∼ 𝒩 (**0**, *I*) *and a family of n* × *n matrices* 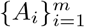 *satisfy*

1. 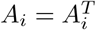 *for i* = 1, …, *m*,
2. rank(*A*_*i*_) = *r*_*i*_ *for i* = 1, …, *m*,
3. 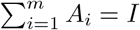

*then each* 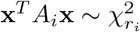 *independently if and only if* 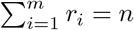.

*Proof*. See Chapter 2 of [48]. □

##### Lemma S1.6

(Null distribution of F-statistic). *Suppose data* **y** ∈ ℝ^*n*^ *is of the form* **y** = *X****β*** + ***ε*** ∈ ℝ^*N*^ *with* ***ε***∼ 𝒩(**0**, *σ*^2^*I*) *and the null hypothesis H****β*** = 0 *holds true, then the F-statistic from Cor. S1.4.1 satisfies*

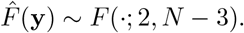

*Proof*. We use the definition o f t he F -statistic in E q. (15). S ince t he p rojection m atrices (*I* − *P*_*u*_) and (*P*_*u*_ − *P*_*r*_) vanish on constant vectors, we may assume without loss of generality that ***µ*** = **0**. Since the projection matrices are idempotent and symmetric, their ranks are given by the dimension of the subspaces onto which they project

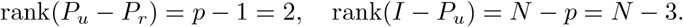

It follows from Theorem S1.5 that the following two quadratic forms have independent chi-squared distributions

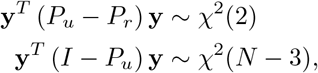

and hence their ratio has an *F* (·; 2, *N* − 3) distribution after scaling by the degrees of freedom. □

##### Lemma S1.7.

*Given projection matrices P*_*A*_ *and P*_*B*_ *that satisfy P*_*A*_*P*_*B*_ = 0 *and* **x** ∼ 𝒩 (***µ***, *σ*^2^*I*), *the random variables* 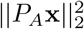 *and* 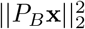 *are independent*.

*Proof*. Since *P*_*A*_ and *P*_*B*_ are commuting projectors they can be simultaneously diagonalized. Working in their common eigenbasis, their characteristic function 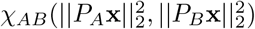 clear^2^ly satisfies

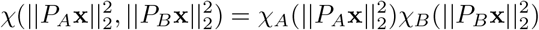

which implies independence.

##### Lemma S1.8

(Alternative distribution of F-statistic). *Suppose data* **y** ∈ ℝ^*N*^ *is of the form* **y** = *X****β*** + *ε* ∈ℝ^*N*^ *with* ***ε*** ∼ 𝒩 (**0**, *σ*^2^*I*) *and the alternative hypothesis H****β*** ≠ 0 *holds true, then the F-statistic from Cor. S1.4.1 has a noncentral F-distribution*

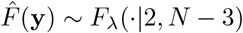

*with noncentrality parameter* 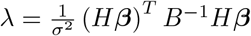 *in whichB = H(X*^*T*^*X)*^−1^*H*^*T*^.

*Proof*. The vector appearing in the numerator of the F-statistic in Eq. (15) satisfies

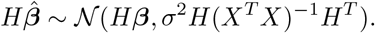

Since *X* and *H* satisfy the non-degeneracy conditions of Lemmas S1.1-S1.3, *B* is symmetric positive definite and therefore has a Cholesky factorization. Let *B*^−1*/*2^ be a Cholesky factor of *B*^−1^, and since 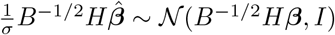,the numerator of Eq. (15) can be rescaled to have a noncentral chi-squared distribution

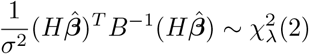

with noncentrality parameter 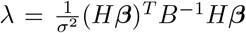.Since (*I* − *P*_u_)*X****β*** = 0 for any ***β*** ∈ ℝ^*p*^, the denominator of Eq. (15) satisfies

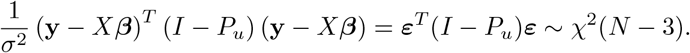

Finally Lemma S1.7 ensures that the numerator and denominator of Eq. (15) are independent and the claim follows.

**Proof of Theorem 3.1**. The power *γ* of a hypothesis test is given by

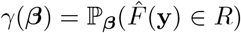

in which *R* ⊂ ℝ^*N*^ is the rejection region, **y** ∈ ℝ^*N*^ is the data and 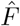 is the F-statistic. Lemma S1.6 implies 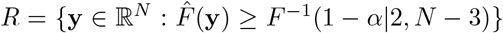 and we may conclude from Lemma S1.8 that ℙ_***β***_(*x ≥ c*) = 1 − *F*_*λ*_(*c*; 2, *N* − 3) for any *c >* 0. Hence we have

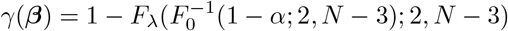

with 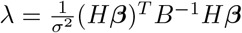.

#### S1.2 Monotonicity of the noncentral F-distribution

Throughout our work, we treat power maximziation and maximization of the noncentrality parameter as equivalent optimization problems. This equivalence is justified because the F-distribution is a monotone function of its noncentrality parameter.

##### Proposition S1.1

(Monotonicity of the noncentral F distribution). *The noncentral F distribution F*_*λ*_(*x*; *n*_1_, *n*_2_) *with degrees of freedom n*_1_ = 2 *and n*_2_ = *N*− 3 *>* 0 *is a monotone decreasing function of the noncentrality parameter λ*.

We prove Prop. S1.1 by expressing the F-distribution as an infinite series, justifying term-wise differentiation, and verifying that the derivative of the distribution with respect to its noncentrality parameter has a definite sign.

##### Lemma S1.9

(Series representation of noncentral F-distribution). *The cumulative distribution function F* (*x*; *n*_1_, *n*_2_, *λ*) *for a noncentral F-distributed random variable with degrees of freedom n*_1_, *n*_2_ *and noncentrality parameter λ can be expressed as a series*

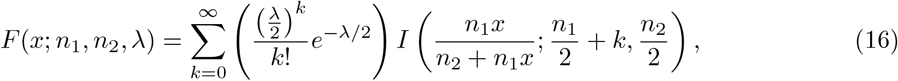

*in which I*(*z*; *a, b*) *is the regularized incomplete beta function, defined by*

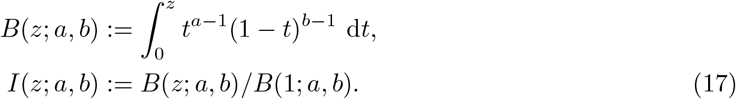

*Proof*. A derivation of this formula is given in [49]. □

##### Lemma S1.10.

*For z* ∈ [0, 1] *and a >* 0, *b >* 0, *the regularized incomplete beta functions obey*

- *I*(*z*; *a* + 1, *b*) ≤ *I*(*z, a, b*),
- lim_*a*→∞_ *I*(*z*; *a* + 1, *b*)*/I*(*z*; *a, b*) = *z*.

*Proof*. The coefficient *B*(1, *a, b*) in Eq. (17) can be expressed as

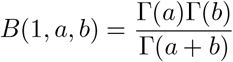

in which 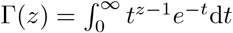 is the gamma function [50]. Use integration by parts and the identity Γ(*z* + 1) = *z*Γ(*z*) to obtain

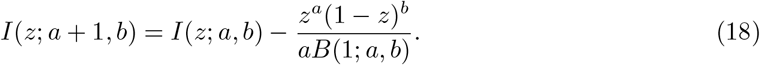

Since *z* ∈ [0, 1] the final term in Eq. (18) is of definite sign, hence

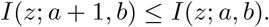

To obtain the asymptotic result, rearrange Eq. (18) and integrate by parts to obtain

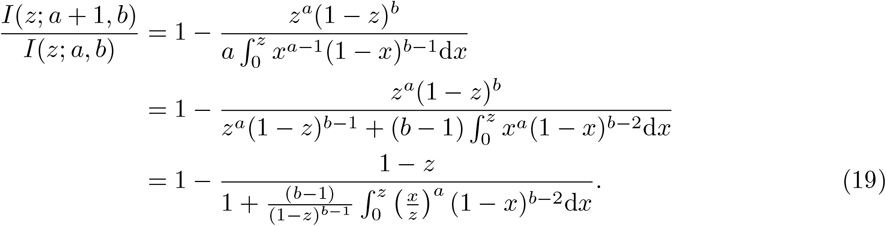

Since *b >* 0, *x < z* and *z* ∈ [0, 1], we have

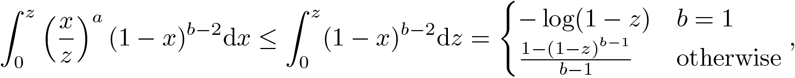

and hence we can evaluate the desired limit using the dominated convergence theorem to obtain

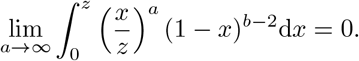

So we conclude from Eq. (19) that lim_*a*→∞_ *I*(*z*; *a* + 1, *b*)*/I*(*z*; *a, b*) = *z*.

##### Lemma S1.11.

*Let T*_*k*_(*z, n*_1_, *n*_2_, *λ*) *be the terms of the series expansion in Eq*. (16) *in which* 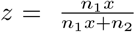 *and λ >* 0. *The terms T*_*k*_(*z, n*_1_, *n*_2_, *λ*) *satisfy the following:*

- 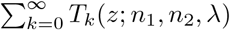 *converges*,
- 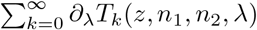 *converges uniformly for all λ >* 0.

*Proof*. Convergence of 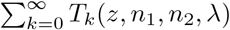 follows from the ratio test

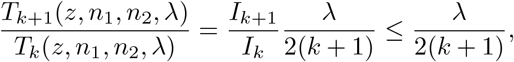

which implies lim 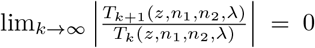 for any *λ >* 0. Uniform convergence of the derivatives can be verified using the Weierstrass M-test. To construct a majorant for *T*_*k*_(*z, n*_1_, *n*_2_, *λ*), notice that ∂_*λ*_*T*_*k*_ has critical points at *λ* = 2*k* and *λ* = 0. We know that *T*_*k*_(*z, n*_1_, *n*_2_, 0) = 0 and lim_*λ*→∞_ *T*_*k*_(*z, n*_1_, *n*_2_, *λ*) = 0 so

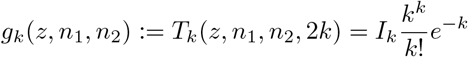

must be the global maximum of *T*_*k*_(*z, n*_1_, *n*_2_, *λ*) on the interval *λ* ∈ [0, ∞). This majorant also leads to a bound on the derivative, since for *k* = 0, we have 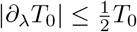 and for *k >* 0

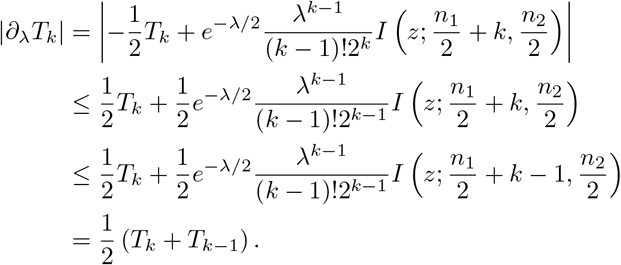

To prove uniform convergence of ∑ _*k*_ ∂_*λ*_ *T*_*k*_, it remains to show that 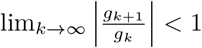.Indeed, we have

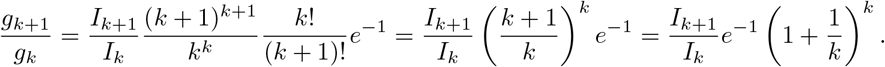

We know 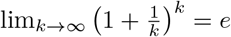 and so

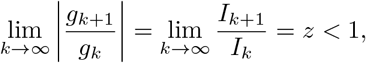

with the last equality justified by the second claim of Prop. S1.10.

**Proof of Proposition S1.1**. Use Prop. S1.9 to express *F* (*x*; *n*_1_, *n*_2_, *λ*) as

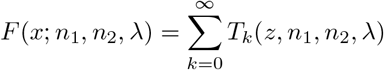

in which 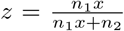.Since *n*_2_ = *N* −3 *>* 0 by assumption, it follows that *z*∈ [0, 1] and we may apply Lemma S1.10 and Lemma S1.11 to justify term-wise differentiation of the series. Differentiate term-wise and apply the same reasoning as in the proof of Lemma S1.11 to find

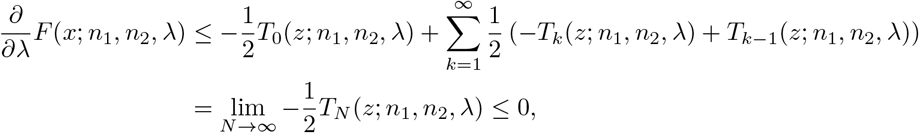

which verifies that *F* (*x*; *n*_1_, *n*_2_, *λ*) is monotone decreasing for all *λ ≥* 0.

#### S1.3 Worst-case power maximization

We prove Theorem 3.2 in two steps. We first show that the worst-case value of the noncentrality parameter is related to an eigenvalue of the matrix *B*(**t**; *f*) defined in Eq. (6). We then compute this eigenvalue for equispaced designs and show that it is optimal using a classical result from optimal experimental design.

##### Lemma S1.12.

*Let H be the hypothesis matrix defined in Eq*. (4) *and let B* = *B*(**t**; *f*) *be the matrix given in Eq*. (6). *The lowest value of the noncentrality parameter across all signals of a given amplitude and frequency*

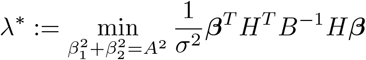

*is given by*

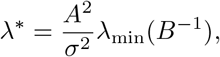

*in which λ*_min_(·) *is the smallest eigenvalue of a symmetric matrix*.

*Proof*. Start by simplifying the definition of *λ*^*^ as

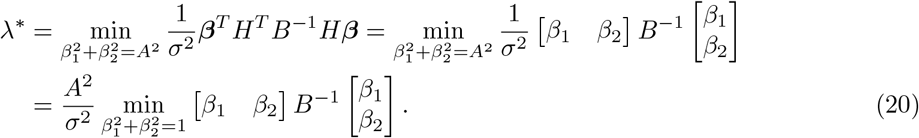

We can set 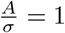 without loss of generality since it appears homogeneously in Eq. (20). Lemma S1.2 ensures that (*X*^*T*^ *X*)^−1^ is symmetric positive definite when *X* is the design matrix for the cosinor model. Since the rhythm detection hypothesis matrix has full row rank, Lemma S1.3 ensures that *B* is symmetric positive definite. It follows that *B*^−1^ has a Cholesky factorization *B*^−1^ = *LL*^*T*^ for some *L* ∈ ℝ^2*×*2^. We may rewrite Eq. (20) in terms of *L* and obtain

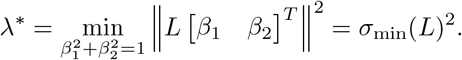

The last equality is justified by the max-min principle for singular values

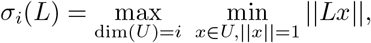

with *σ*_2_ ≤ *σ*_1_, so *i* = 2 gives the smallest singular value. The result follows by recalling that *B*^−1^ has real eigenvalues and thus *σ*_min_(*L*)^2^ = *λ*_min_(*B*^−1^). □

The next two lemmas allow are used in computing the minimum eigenvalue of *B*(**t**; *f*)^−1^ for equispaced designs.

##### Lemma S1.13.

*Let B* = *B*(**t**; *f*) *be the matrix B* = *H*(*X*^*T*^ *X*)^−1^*H*^*T*^, *in which X* = *X*(**t**; *f*) *is the design matrix for the one-frequency cosinor model and H is the hypothesis matrix. The inverse of B can be expressed as*

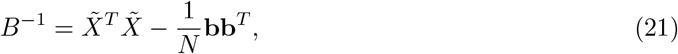

*in which* 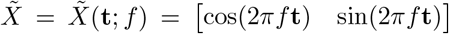 *is the design matrix for the mean-free cosinor model Y* (*t*) = *β*_1_ cos(2*πft*) + *β*_2_ sin(2*πft*) + *ε*(*t*) *and* **b** *is the vector of time-averages*

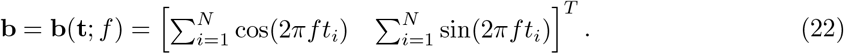

*Proof*. Compute the inverse of *X*^*T*^ *X* using the Schur complement formula and act with *H* to find that the only remaining term agrees with the right hand side of Eq. (21).

##### Lemma S1.14.

*If* **t** ∈ ℝ^*N*^ *is equispaced over a cycle of frequency f and N ≥* 3 *then*

1. 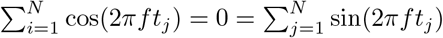
2. *X*(**t**; *f*)^*T*^ *X*(**t**; *f*) = diag(*N, N/*2, *N/*2).

*Proof*. Since the measurements are equispaced over a cycle of frequency *f*, we may assume without loss of generality that they are located at the roots of unity 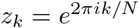 for *k* = 0, …, *N* − 1. The time averages can be evaluated using a finite geometric series to find

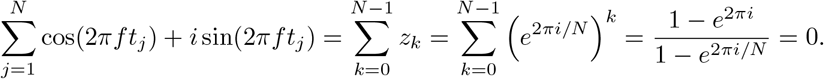

The same argument applied to the entries of the *X*^*T*^ *X* reveals that *X*^*T*^ *X* = diag(*N, N/*2, *N/*2). □

It follows from Lemma S1.14 that any equispaced **t** ∈ ℝ^*N*^ with *N >* 3 will produce a design matrix that satisfies

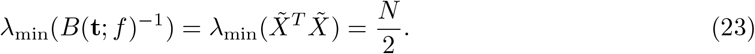

We verify that the value *N/*2 on the right hand side of Eq. (23) is globally optimal by using an equivalence theorem from optimal experimental design. These classical theorems are reviewed in [51]. For our purposes, it is convenient to use a more modern statement of the result, which we adapt from [21].

##### Theorem S1.15

(Elfving optimality condition, [21, Theorem 7.22]). *Let* Ξ *be the design space*

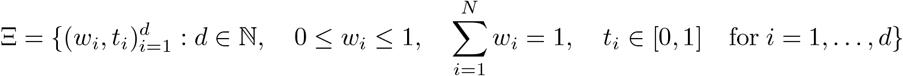

*and let* ℳ (Ξ) *be the corresponding space of moment matrices*

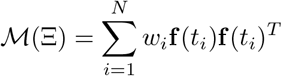

*in which* **f** (*t*) = cos(2*πft*) sin(2*πft*) ^*T*^ *is the mean-free cosinor model. A moment matrix M*^*^ ∈ ℳ (Ξ) *is Elfving optimal in the sense that*

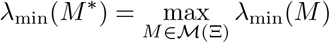

*if and only if there exists a positive semidefinite matrix E* ∈ ℝ^2*×*2^ *such that* tr (*E*) = 1 *and*

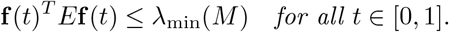

**Proof of Theorem 3.2**. Power and noncentrality parameter maximization for a given acrophase, frequency, and amplitude are equivalent by Prop. S1.1. Maximizing the noncentrality parameter across all acrophases is equivalent to maximizing the minimum eigenvalue of *B*(**t**; *f*)^−1^ by Lemma S1.12. It remains to show that equispaced designs with *N >* 3 measurements provide optimal solutions to the eigenvalue programming problem and that their power is acrophase-independent.

First use Lemma S1.13 to rewrite the eigenvalue problem as

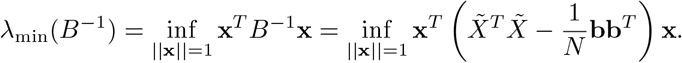

We clearly have 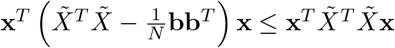 for any **x** ∈ ℝ^2^, and consequently

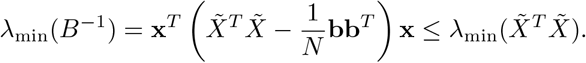

If **t** is equispaced, then **b** = **0** by Lemma S1.14 and so 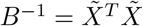 and we find

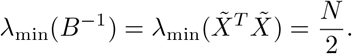

To apply Theorem S1.15 and complete the verification of global optimality, notice that every equispaced design **t** corresponds to a standardized design 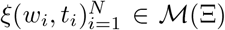 with *w*_*i*_ = 1*/N* with design matrix *M* (*ξ*) = diag(1*/*2, 1*/*2). Notice that we have

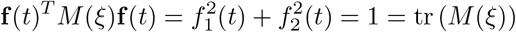

which verifies global optimality by Theorem S1.15. Finally, explicit computation confirms that the noncentrality of an equispaced design satisfies

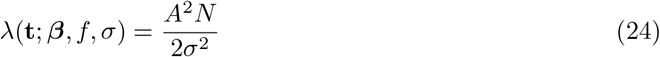

where 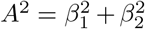 is the amplitude. Notice that Eq. (24) does not depend on the acrophase of the signal, as required.

#### S1.4 Power analysis at the Nyquist rate

For a sample size *N* and equispaced design **t**_*N*_, power analysis at the Nyquist rate 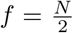 requires some additional consideration. Even if the sample size satisfies *N >* 3, the measurements will not cover sufficiently many distinct phases for the design matrix to be full rank. Hence, the methods developed so far in this section are not immediately applicable for computing the worst-case power of an equispaced design at the Nyquist rate. The following proposition allows us to circumvent this issue.

##### Proposition S1.2

(Noncentrality parameter at the Nyquist rate). *For N >* 3, *the noncentrality parameter of an equispaced design* **t**_*N*_ *satisfies*

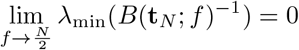

*and hence the worst-case power is given by γ* = *α, in which α is the type-I error rate*.

*Proof*. Evaluate the entries of 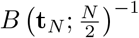 using Eq. (21) and a finite geometric series to obtain

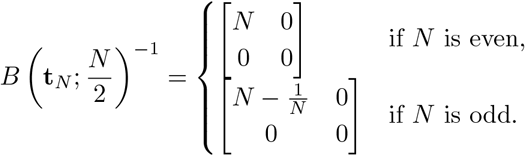

Since eigenvalues are continuous with respect to entrywise matrix convergence (see for instance [52, Proposition 2.4.9.2]), it follows that

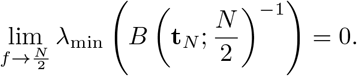

When the noncentrality parameter *λ* = 0, the power expression simplifies to

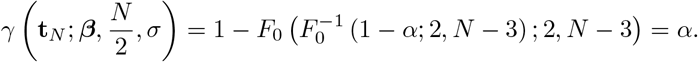

#### S1.5 Unions of equispaced designs

Consider a family of equispaced designs 𝒯= {**t**^(1)^, …, **t**^(*K*)^} with each 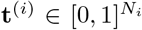 and suppose the design matrix of each individual equispaced **t**^(*i*)^ satisfies the non-degeneracy condition from Lemma S1.1. Our goal here is to show that the union of these designs

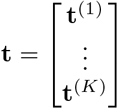

retains the optimality properties of the individual designs. The key observation comes from Lemma S1.13 and noticing that the time-average vector **b**(**t**; *f*) defined in Eq. (22) vanishes since

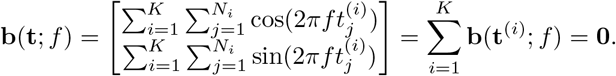

Hence, using Eq. (21), matrix *B*(**t**, *f*) = *H (X*(**t**; *f*)^*T*^ *X*(**t**; *f*))^−1^ *H*^*T*^ simplifies to

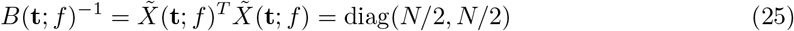

in which 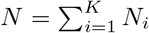.It follows from Eq. (25) that the phase-independent power and optimality properties of the individual designs are still applicable in the full design **t**.

### S2 Supplementary figures

**Supplementary Figure 1:**
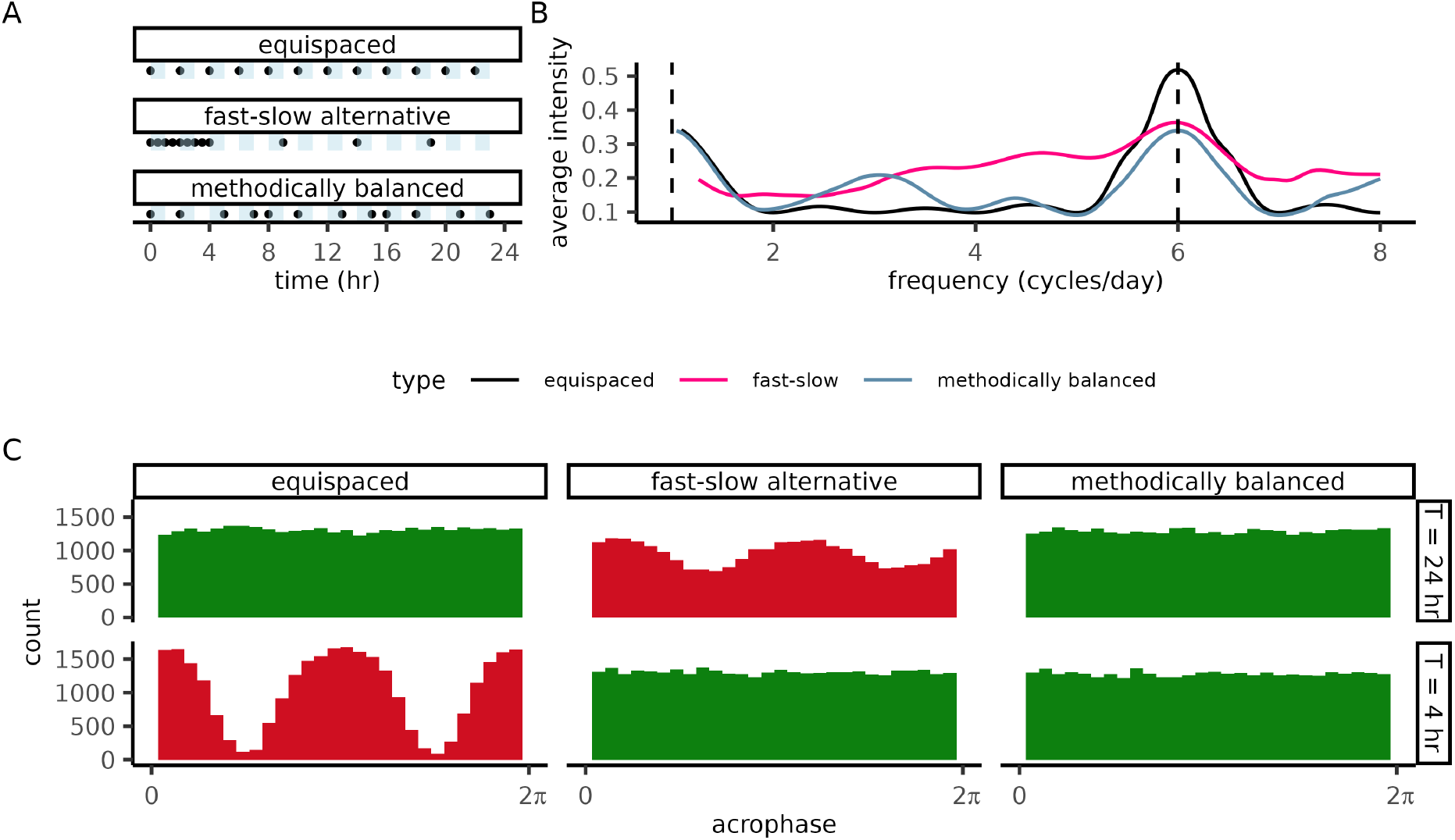
An irregular design detects 24hr and 4hr rhythms at all acrophases. Each oscillator (*n* = 10^5^) in the simulated dataset was assigned a 24hr or 4hr period and a uniformly random acrophase *ϕ* ∈ [0, 2*π*). Measurements are simulated with Gaussian white noise at each measurement time. **(A)** Measurement schedules for (top) a traditional equispaced design, a (middle) fast-slow irregular design, and (bottom) a methodically constructed irregular design. The shaded bars represent 2hr increments and dots indicate sample collection (*N* = 12 samples for each design). **(B)** The average intensity of a Lomb-Scargle periodogram for each design. The true periods in the system are marked by the dashed vertical lines. **(C)** True acrophases of statistically significant oscillators (*p <* 0.05) detected by cosinor analysis at each of the true periods. Distributions with phase-dependent detection are shown in red to emphasize that the distribution’s variability is due to a statistical artifact. Simulation parameters: amplitude 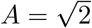,noise strength *σ* = 1, acrophase *ϕ*∼ Unif(0, 2*π*), period *T* = 4hr, 24hr. The methodically-constructed design was generated using mixed-integer conic programming in PowerCHORD.

**Supplementary Figure 2:**
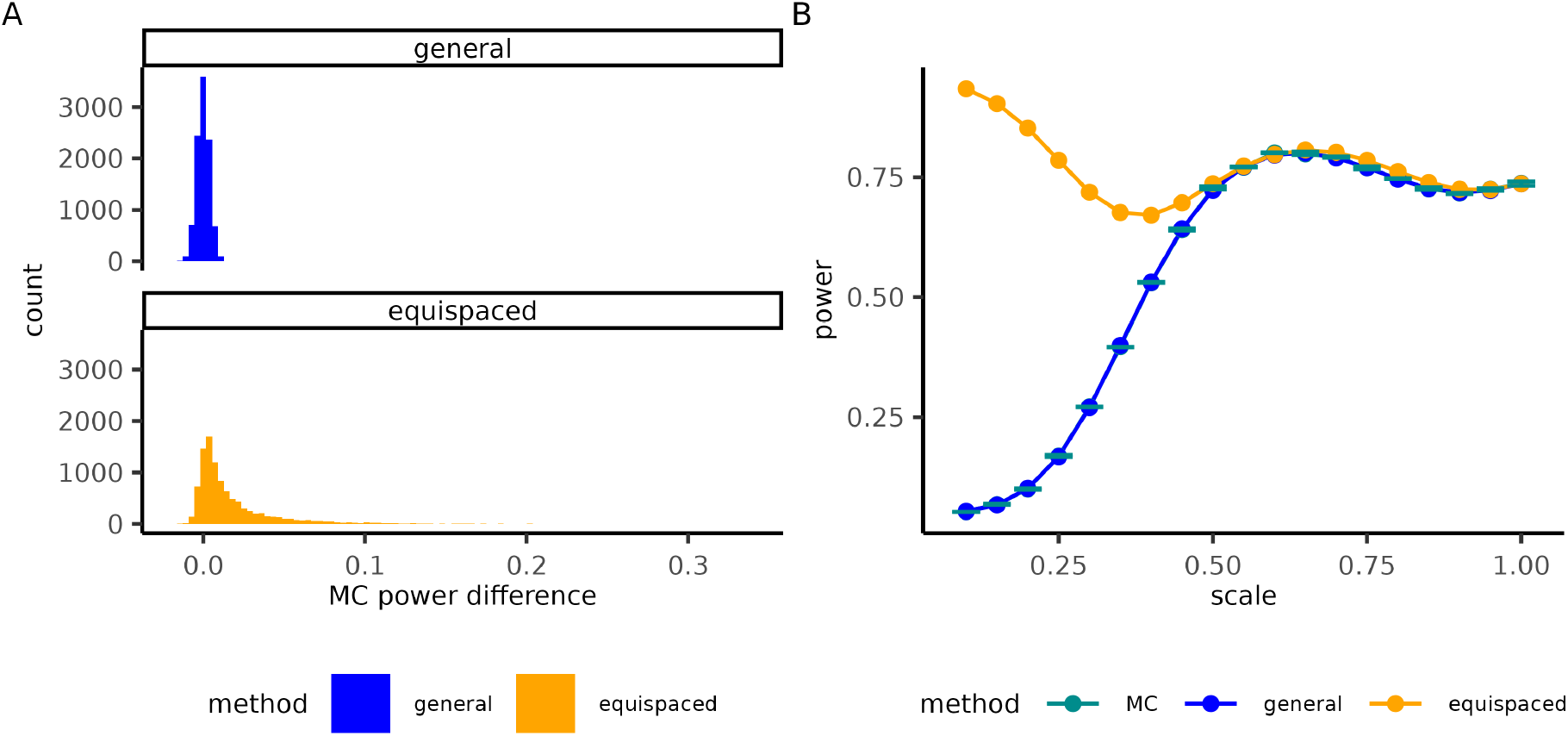
The general power formula is necessary for accurate power analysis. **(A)** Comparison of our power formula and the equispaced formula to Monte Carlo estimates for randomly generated designs in which *t* ∼ unif([0, 1]) for each measurement time *t*. On average, the equispaced formula tended to over-estimate the power of such designs. Parameters: sample size *N* = 8, amplitude *A* = 2, frequency *f* = 1, acrophase *ϕ* = *π*, noise strength *σ* = 1. **(B)** We computed the power of designs **t**_*N,κ*_ = *κ***t**_*N*_, where **t**_*N*_ is an *N* measurement equispaced design and *κ >* 0 is a scale factor. As *κ* shrinks, the irregularity in the design becomes more pronounced and the equispaced formula diverges from the Monte Carlo power estimates. Parameters: *N* = 24, *A* = 1, *f* = 1, *ϕ* = 0

**Supplementary Figure 3:**
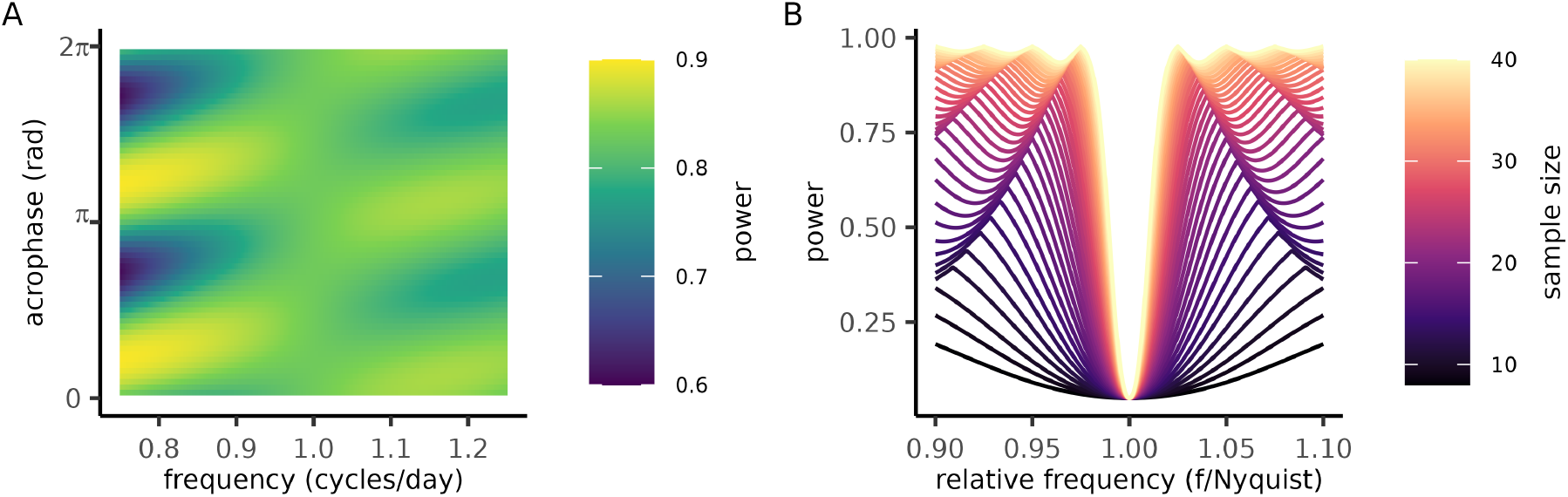
Phase-dependence of equispaced power near critical frequencies. **(A)**The power (color) of an equispaced design (*N* = 24 samples) evaluated at each frequency (x-axis) and acrophase (y-axis). The power is independent of phase when the frequency reaches *f* = 1 because the design is equiphase at this frequency. **(B)** The worst-case power of equispaced designs (sample size 8 ≤ *N* ≤ 40) as a function of frequency, with frequency scaled relative to Nyquist rate of each design (*f*_rel_ = *f/f*_Nyq_). Parameters: amplitude *A* = 1, noise strength *σ* = 1.

**Supplementary Figure 4:**
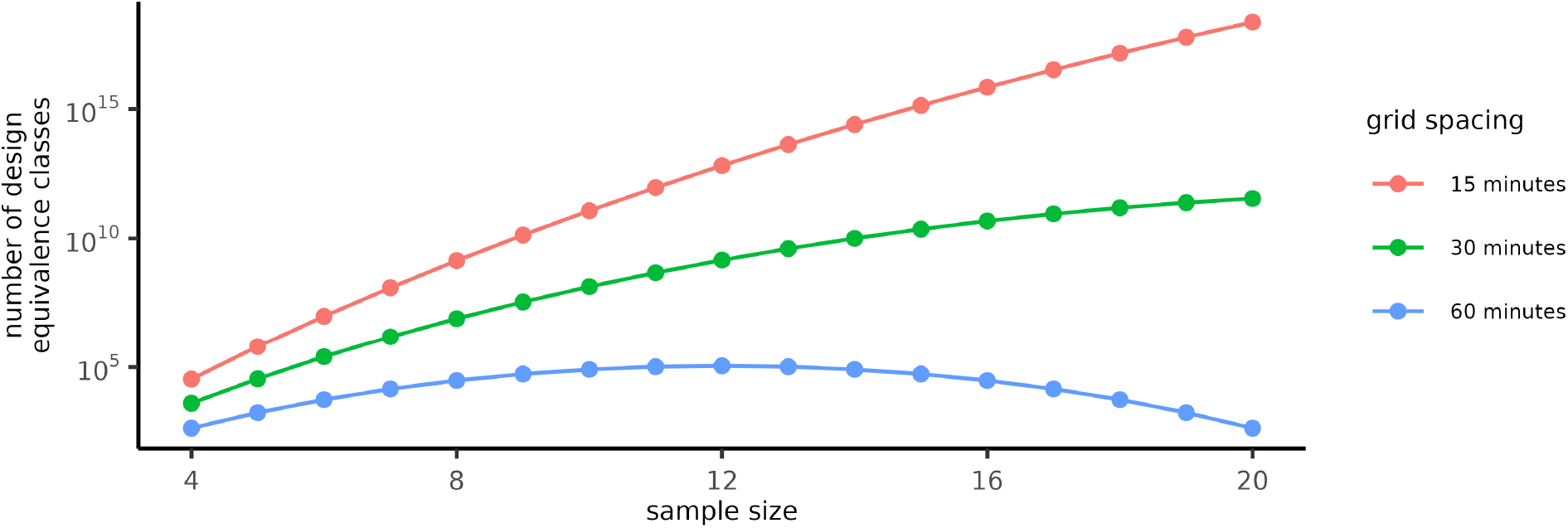
The number of experimental design equivalence classes grows rapidly with sample size. For a given sample size (x-axis) and grid spacing (color), the number of design equivalence classes (y-axis) can be calculated using Eq. (10). Designs are in the same equivalence class if they can be transformed into one another by a cyclic shift (i.e. *t* → *t* + *k/N*_*t*_ mod 1, for some 1 ≤ *k* ≤ *N*_*t*_ assuming measurements are in the interval [0, 1] and confined to a grid of spacing 1*/N*_*t*_).

**Supplementary Figure 5:**
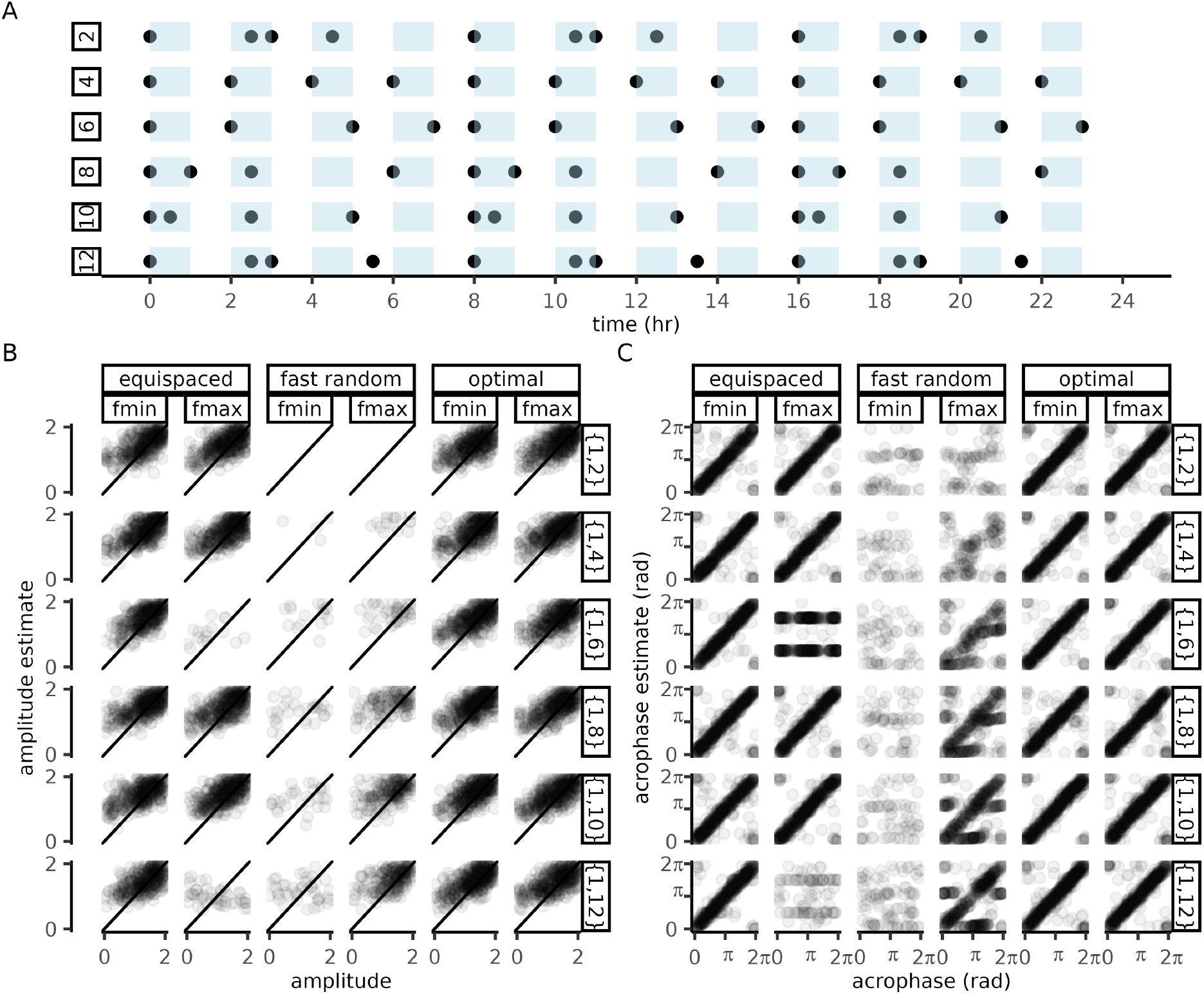
Bifrequency optimal designs exhibit low bias at both intended frequencies. We computed bifrequency optimal designs for frequency priors *ν* = (1, *f*) with *f* ∈(2, 4, 6, 8, 10, 12) and a sample size *N* = 12. **(A)** Repetitive patterns appear in the measurement times of the bifrequency optimal designs. **(B)** Each design was tested for bias by comparing the true values of the amplitude and acrophase to their cosinor estimates after filtering for statistically significance. With the exception of signals at integer multiples of the Nyquist rate, equispaced and optimal designs performed similarly. At the Nyquist multiples, the optimal designs exhibited much less bias than equispaced designs. A randomly generated design (*t*∼ unif(0, 1*/*12)) with measurements confined to a short timescale was included as a reference. The random design performed poorly at low frequencies and improved as the higher frequency approaches the scale on which its points are distributed.

**Supplementary Figure 6:**
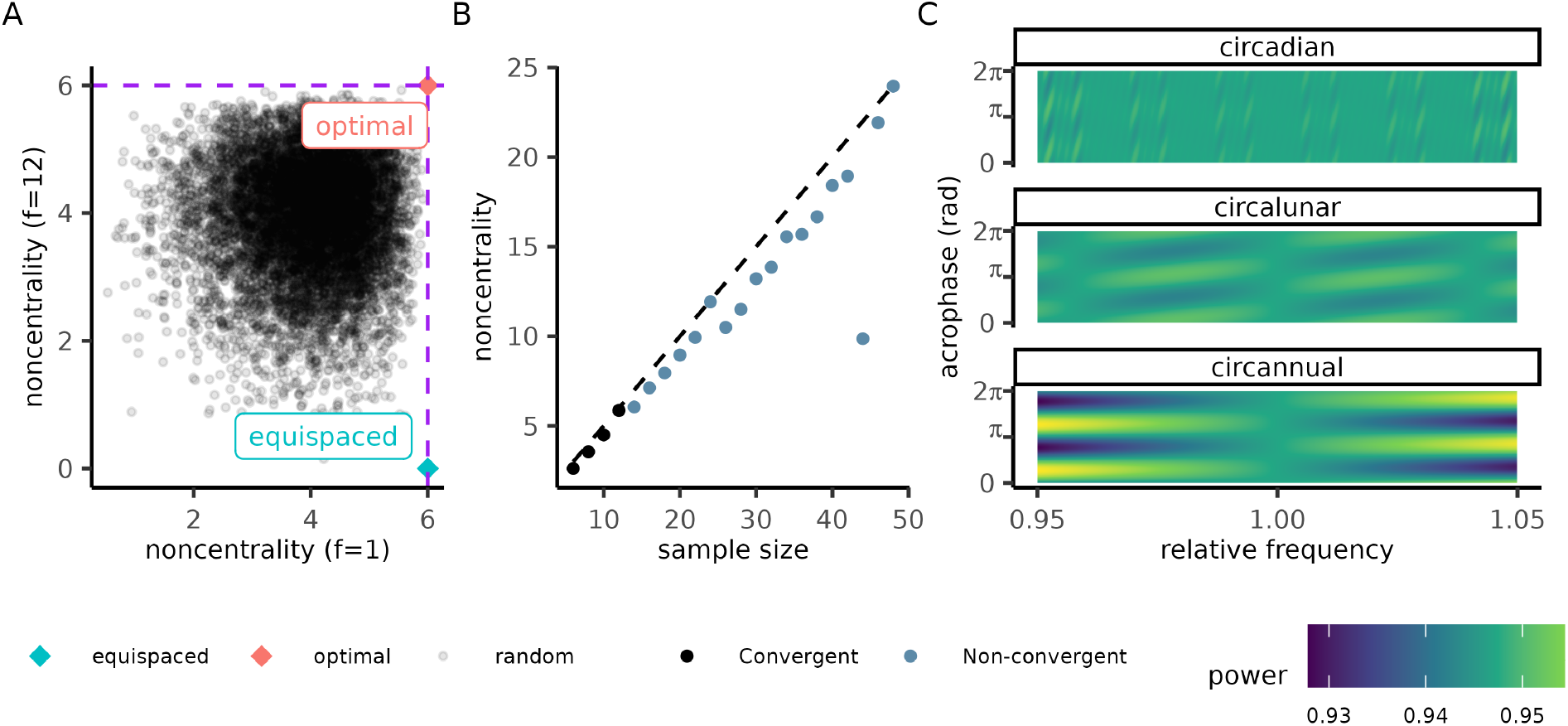
Bifrequency and trifrequency optimal designs at critical frequencies. **(A)** Non-centrality parameters for the two periodicities of interest (f=1 and f=12) for the optimal design (red dot) an equispaced design (blue dot) and an ensemble (*n* = 10^4^) of randomly generated designs. The theoretical maximum value of the non-centrality parameter (*λ* = *N/*2; Theorem 3.2) is indicated by dashed lines. **(B)** Designs were generated to maximize power at the first *N/*2 harmonics (*f* ∈ (1, …, *N/*2)) for each sample size *N* (x-axis). The performance of each design is summarized by the lowest value of its non-centrality parameter (y-axis) across all harmonics included in the optimization. Color indicates convergence of the conic program within 1hr of computation time. The optimal noncentrality parameter in a single frequency design for each sample size (*λ* = *N/*2) is shown for reference (dashed line). For sample sizes 1≤ *N <* 12 measurements were confined to a 36 point grid, for 12≤ *N <* 24 a 48 point grid, and for 24 *< N* a 96 point grid. **(C)** The trifrequency optimal design with all measurements confined to the first month achieves phase independent power at 24 hr (circadian),28 day (circalunar), and 12 × 28 = 336 day (circannual) periods. Parameters: Amplitude 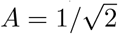.

## Notes

https://github.com/t-silverthorne/PowerCHORD

### Competing Interest Statement

The authors have declared no competing interest.

### Summary of Updates

The power formula in Section 3.1 has been generalised to hold for arbitrary measurement times. Two additional optimisation methods (brute-force searching and conic programming) have been added to the repository and studied in Sections 3.3-3.4.

